# Transformation of Primary Sensory Cortical Representations from Layer 4 to Layer 2

**DOI:** 10.1101/2021.09.17.460780

**Authors:** Bettina Voelcker, Simon Peron

**Affiliations:** Center for Neural Science, New York University, 4 Washington Place Rm. 621 New York, NY 10003

## Abstract

Sensory input arrives from thalamus in cortical layer (L) 4, from which it flows predominantly to superficial layers, so that L4 to L2 constitutes one of the earliest cortical feedforward networks. Despite extensive study, the transformation performed by this network remains poorly understood. We use two-photon calcium imaging in L2-4 of primary vibrissal somatosensory cortex (vS1) to record neural activity as mice perform an object localization task with two whiskers. We find that touch responses sparsen but become more reliable from L4 to L2, with superficial neurons responding to a broader range of touches. Decoding of sensory features either improves from L4 to L2 or remains unchanged. Pairwise correlations increase superficially, with L2/3 containing ensembles of mostly broadly tuned neurons responding robustly to touch. Thus, from L4 to L2, cortex transitions from a dense probabilistic code to a sparse and robust ensemble-based code that improves stimulus decoding, facilitating perception.

## INTRODUCTION

Primary sensory cortices are organized into columns processing similar stimulus features and exhibiting precise interlaminar wiring (Mountcastle 1997, Narayanan et al 2017). Canonically, thalamic input mostly targets layer (L) 4, with weaker projections targeting L3 and L5 (Constantinople & Bruno 2013, Ji et al 2016, Meyer et al 2010, Wimmer et al 2010). The strongest interlaminar projection from L4 is to L3, and from L3, to L2 (Hooks et al 2011, Lefort et al 2009), making primary sensory L4 to L2 one of the earliest cortical feedforward networks. Despite extensive study, the computational role of this feedforward network remains unclear (Adesnik & Naka 2018, Douglas & Martin 2004, Petersen & Crochet 2013).

Progressive reduction in the fraction of neurons responding to a particular stimulus, or sparsification, has been proposed as a core function of feedforward sensory processing. By producing a reliable and robust representation of the external world, sparsification can facilitate perceptual readout (Barlow 1972, Olshausen & Field 2004, Wolfe et al 2010). Sparsification is often accompanied by the emergence of neurons with highly selective responses that pool across a particular dimension of the stimulus. For instance, in the visual dorsal stream, receptive fields expand in angular extent and become increasingly object-selective (Kobatake & Tanaka 1994, Reddy & Kanwisher 2006). Sparsification from L4 to L2 has been observed in primary sensory cortices (Niell & Stryker 2008, O’Connor et al 2010b, Sakata & Harris 2009), as have broader receptive fields (Hirsch & Martinez 2006, Simons 1978, Winkowski & Kanold 2013). Because sparsification and receptive field broadening are typically studied in isolation, it remains unclear whether the transition to broader receptive fields coincides with sparsification. Moreover, most studies of receptive field expansion were performed in anesthetized animals, which can alter receptive fields: in the vibrissal system, thalamic and cortical neurons become responsive to a larger number of whiskers under anesthesia (Friedberg et al 1999, Simons et al 1992). Finally, it is unclear whether sparsification and receptive field expansion improve stimulus decoding, as has been proposed (Barlow 1972, Olshausen & Field 2004, Wolfe et al 2010).

In addition to sparse representations and expanded receptive fields, layer 2/3 contains groups of correlated neurons tuned to similar features, or ensembles (Buzsaki 2010, Carrillo-Reid et al 2017, Harris & Mrsic-Flogel 2013, Hebb 1949). In mouse L2/3, neurons with correlated activity are more likely to be directly connected (Cossell et al 2015, Lee et al 2016), enabling computations such as pattern completion (Carrillo-Reid et al 2019, Marshel et al 2019) and amplification (Peron et al 2020). Because synchronous L2/3 activity triggers strong feedback inhibition (Mateo et al 2011), ensembles have been proposed as contributing to sparseness by evoking such inhibition (Barth & Poulet 2012). Furthermore, due to their capacity for pattern completion and amplification, ensembles should produce a more reliable neural code (Buzsaki 2010, Harris & Mrsic-Flogel 2013). Due to the dense sampling needed for studying sparse populations and ensembles, however, it remains unclear whether the majority of the sparse, stimulus responsive neurons in L2/3 belong to ensembles. If the initial stages of cortical sensory processing implement a transition to an ensemble-based code, this would argue for a central role of ensembles in cortical function (Hebb 1949, Sakurai 1999).

We examine the transformation in vS1 touch representations from L4 to L2. In mouse vS1, thalamic input from individual whiskers projects predominantly to small, ~300 μm diameter patches of cortex known as ‘barrels’ (Woolsey & Van der Loos 1970). The circuitry in vS1 L2/3 is relatively well characterized (Petersen & Crochet 2013), making it an ideal system for the study of superficial layers. To achieve dense sampling (Peron et al 2015a), we employ volumetric calcium imaging (Peron et al 2015b) in transgenic mice expressing GCaMP6s in excitatory neurons (Daigle et al 2018). Using mice with two spared whiskers performing an object localization task (O’Connor et al 2010a, Peron et al 2015b), we first examine how the distribution of touch neuron receptive fields changes from L4 to L2. Next, we examine robustness and sparseness of touch responses across layers, along with the ability of neurons to decode touch features. Finally, we examine changes in correlation structure across layers, assessing the role played by groups of co-active neurons, or ensembles, in responding to touch.

## RESULTS

### Mapping dual-whisker barrel cortex responses across layers 2-4

To study how neural representations change from L4 to L2, we trained transgenic mice expressing GCaMP6s in cortical excitatory neurons (Ai162 X Slc17a7-Cre) (Daigle et al 2018) and implanted with a cranial window over vS1 on a two-whisker (typically, C2 and C3; **Fig. 1A; Table S1**) object localization task (O’Connor et al 2010a, Peron et al 2020). Mice reached stable performance (Methods; **Fig. 1B**) in 4.0 ± 2.1 days (mean ± S.D.; n=7 mice). Changes in whisker curvature (Δκ) were used as a proxy for the force impinging on the follicle base and, hence, sensory input (Severson et al 2017) (**Fig. 1C**). We employed continuously distributed pole positions spanning the whisking range, which resulted in four basic types of touch (**Fig. 1D**): whisker 1 protractions (W1P), whisker 1 retractions (W1R), whisker 2 protractions (W2P), and whisker 2 retractions (W2R). The kinematics were consistent across animals (**Fig. 1E**), with certain touch types occurring more frequently than others (**Fig. 1F**).

**Figure 1.**
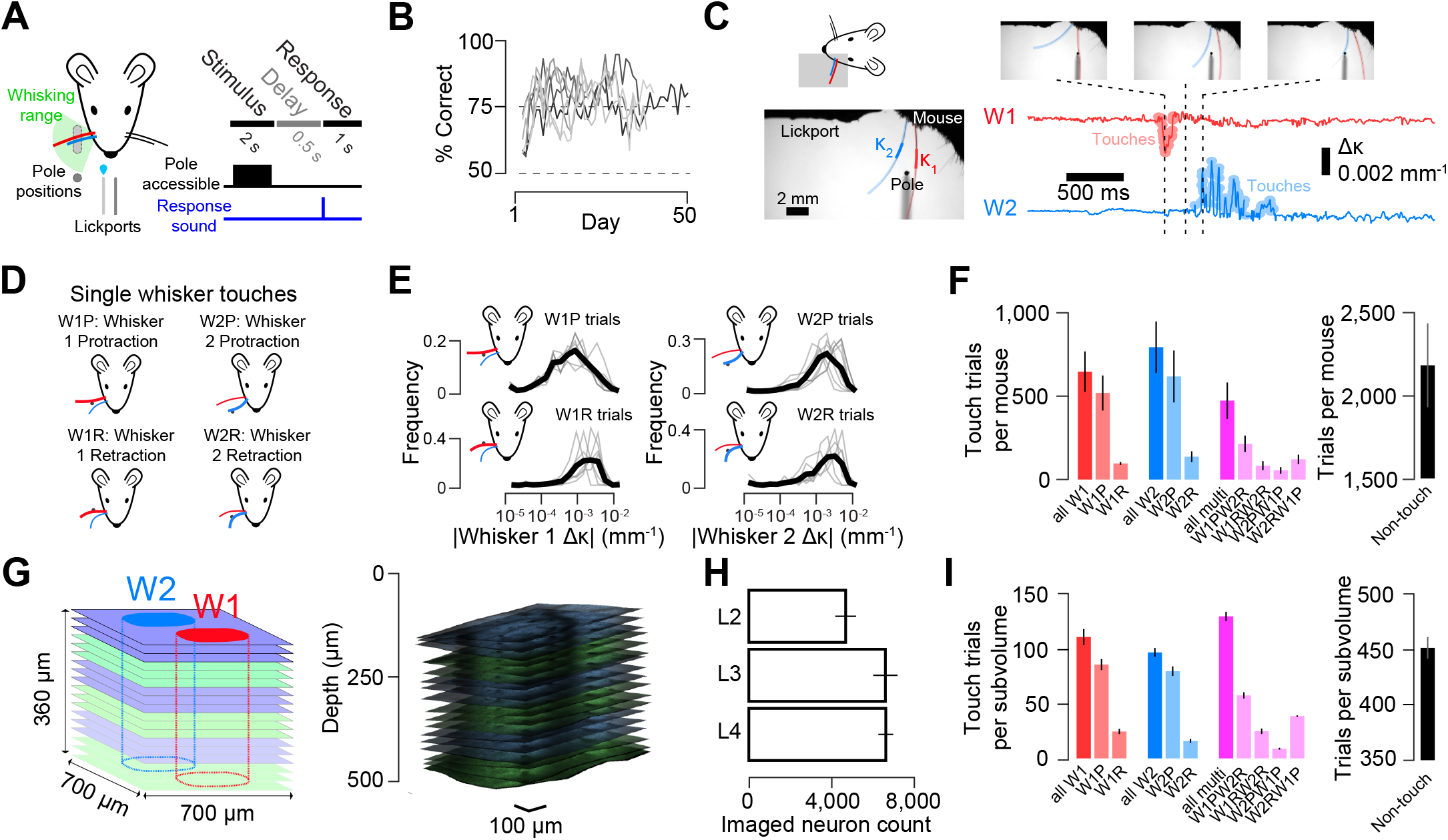
Volumetric calcium imaging during a two-whisker task. **A)** Mice use two whiskers to detect a pole that appears in a proximal position range (light gray) or distal position (dark gray; Methods). Right, task timing. **B)** Training progression (n=7 mice). **C)** Example touch video. Bottom left, the region where curvature is measured is indicated for both whiskers. Bottom right, change in curvature (Δκ) for each of the two whiskers. Moments of touch are highlighted. **D)** Single-whisker touch types. **E)** Distribution of curvature changes for each single-whisker touch type. Dark line, mean across mice (n=7). **F)** Frequency of the trials with a given touch type. Red, whisker 1; blue; whisker 2; magenta, multi-whisker. Bars, mean; line, S.E.M. **G)** Volumetric imaging. Identically colored planes were imaged simultaneously (‘subvolume’ of 3 planes, 20 μm apart; 5-6 subvolumes per mouse). Left, schematic with typical barrel positioning. Right, imaging planes from example mouse plotted after alignment to reference stack (Methods). **H)** Neuron count by layer. Bars, mean across animals; lines, S.E.M **I)** Number of trials per subvolume (and, hence, neuron) for each touch type for an example animal.

Activity in well-trained mice was recorded for 18.9 ± 4.0 sessions (mean ± S.D.; range: 15 to 23, n=7 mice) using volumetric two-photon calcium imaging (**Fig. 1G**), with 5-7 ‘subvolumes’ of three planes imaged simultaneously (n=38 subvolumes across 7 mice). Each session’s imaging planes were aligned to a per-animal reference stack from which cortical depth and laminar bounds were established (**Fig. S1**). Barrel boundaries were obtained from the neuropil signal (Peron et al 2015b) and visible septa in layer (L) 4 (**Fig. S2**). In L2, L3, and L4, we imaged 3,796 ± 1,076, 5,355 ± 1,264 and 5,372 ± 772 neurons per mouse (n=7 mice), respectively (**Fig. 1H; Table S1**). We obtained 820 ± 285 trials per neuron, distributed across different touch types (**Fig. 1I**).

Individual neurons exhibited diverse touch responses, with different neurons tuned to different combinations of single-whisker touches (**Fig. 2A**). Individual neurons showed increasing responsiveness to stronger touches (Methods; **Fig. 2B**). Response probability increased with increasing touch strength (**Fig. 2C, Fig. S3A**). The fraction of neurons responding to touch also increased with touch strength, as did the aggregate response of the population (**Fig. 2D, Fig. S3B**).

**Figure 2.**
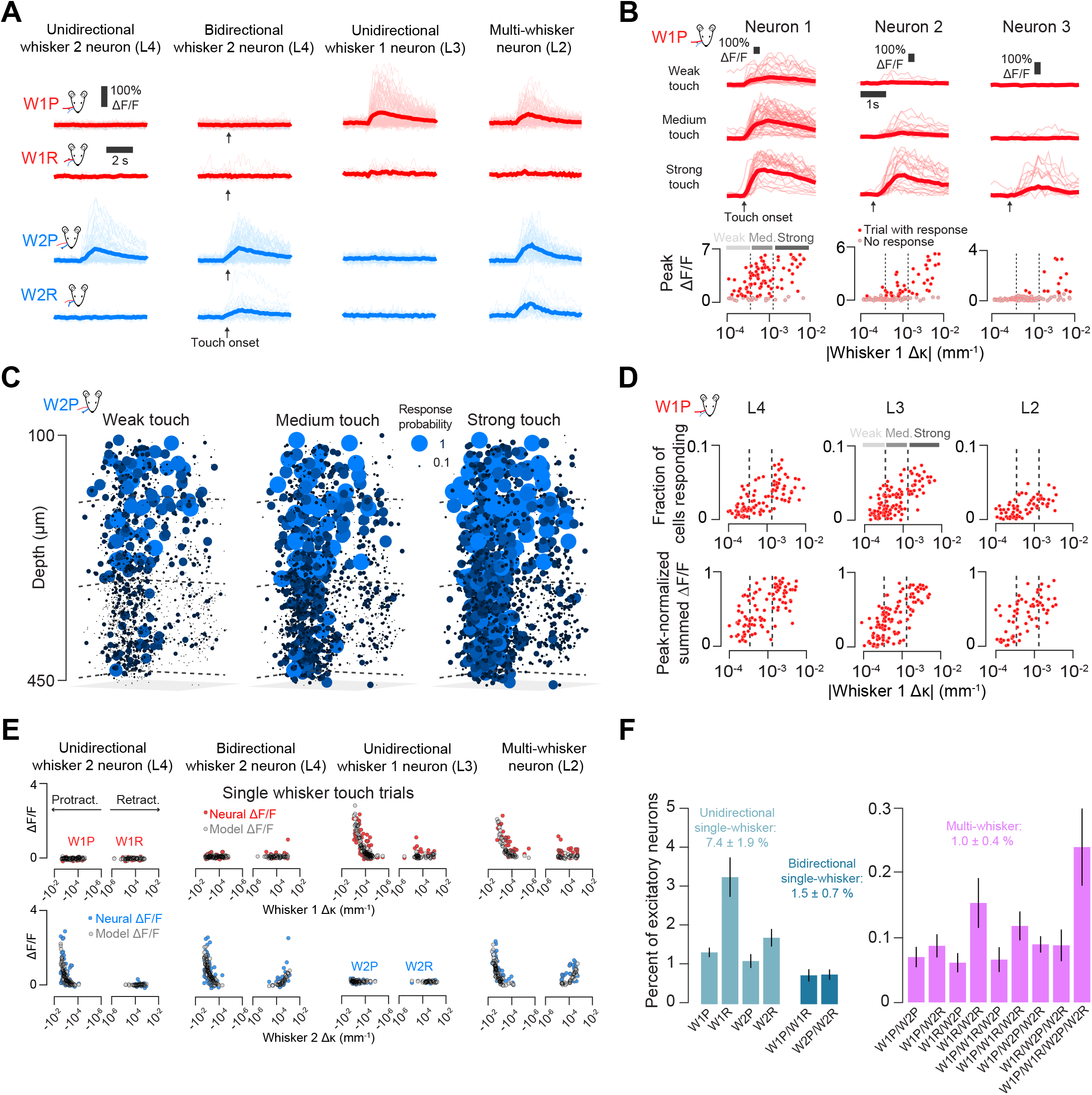
Classification of touch-sensitive neurons. **A)** Example ΔF/F responses to all single-whisker touch types for four neurons. Light color, individual touch-aligned responses; dark color, mean across touches. **B)** Example responses to W1P touch for three neurons. Top, ΔF/F, grouped by touch intensity (weak: bottom third trial Δκ; medium: middle third; strong: top third). Bottom, response as a function of Δκ. Dark red dots, trials with detected response; gray dots, no response. Dotted vertical lines delimit the touch strength bins. **C)** Example response probability plotted for all neurons responding to weak, medium, and strong W2P touch. Ball size and color indicate response probability. **E)** Example population response to W1P touch. Top, fraction of neurons responding as a function of trial ∆κ (Methods). Dots, individual trials. Dashed vertical lines: bins for weak, medium, and strong touch trials. Bottom, normalized summed ∆F/F from the entire responsive pool. **E)** Receptive fields for neurons in **A**. The mean ΔF/F as a function of trial Δκ (Methods) is shown. Red circles, trials where only whisker 1 touched; blue, only whisker 2 touched. Gray circles, model’s predicted ΔF/F. **F)** Frequency of different touch neurons. Frequency is given for each basic touch type combination.

To quantify the sensitivity of neurons to touch, we used an encoding model that predicted neural activity from whisker curvature (**Fig. S4**; Methods). An encoding model score was computed by measuring the Pearson correlation between the predicted and actual ΔF/F (Methods). The model was first fit for each whisker using trials with only single-whisker touches (Methods; **Fig. 2E**). On trials where both whiskers touched, a linear scaling factor was fit and applied to the second contacting whisker’s single-whisker model prediction (**Fig. S4**). Neurons were considered responsive to a given single-whisker touch type if the encoding model score exceeded both 0.1 and an activity-matched temporally shuffled response score for those trials (Methods).

Touch neurons were classified based on the combination of single-whisker touch types they responded to. Across all neurons, 9.9 ± 2.9 % responded to some form of touch. We classified single-whisker touch neurons as unidirectional and bidirectional; neurons that responded on at least one type of single-whisker trial for both whiskers were classified as multi-whisker (Methods; **Fig. 2A**). Across all imaged neurons, 7.4 ± 1.9 % of neurons responded to only one touch direction for one whisker (**Fig. 2F**, ‘unidirectional’ single-whisker neurons), 1.5 ± 0.7 % responded to both directions for a single whisker (‘bidirectional’ single-whisker neurons), and 1.0 ± 0.4 % of neurons responded to both whiskers (‘multi-whisker’).

Multi-whisker touch trials were common (**Fig. 1F**) and exhibited a range of inter-touch intervals (**Fig. S5**). Because the initial model fit only employs touch trials where a single whisker touches the pole (**Fig. S4E**), our fitting approach would miss neurons that responded exclusively to multi-whisker touch. We therefore manually examined neurons showing elevated response probability (Methods) on multi-whisker touches that were not classified as touch neurons. Only two neurons showing exclusive multi-whisker responses were found (**Fig. S5D**). Thus, multi-whisker touch engages neurons that also respond to single-whisker touch.

### Touch receptive fields broaden from L4 to L2

Superficial receptive field broadening has been observed in several primary sensory cortices (Hirsch & Martinez 2006, Simons 1978, Winkowski & Kanold 2013), though past experiments typically employed anesthesia and sampled sparsely. Dense, relatively unbiased sampling with two-photon microscopy allowed us to examine in detail how receptive fields change from L4 to L2 during behavior.

Unidirectional, bidirectional, and multi-whisker neurons showed distinct spatial distributions and varying response probabilities (**Fig. 3A**). ‘Broadly’ tuned neurons – bidirectional single-whisker and multi-whisker neurons – increased in frequency from L4 to L2. Specifically, bidirectional neurons increased in frequency from L4 (0.008 ± 0.006) to L3 (0.021 ± 0.010; L4 vs. L3, p = 0.006), remaining unchanged from L3 to L2 (0.017 ± 0.007; **Fig. 3B**; L3 vs. L2, p = 0.128), as did the relative fraction of multi-whisker neurons (L4: 0.002 ± 0.001, L3: 0.014 ± 0.006, L2: 0.016 ± 0.007; L4 vs. L3, p = 0.001; L3 vs. L2, p = 0.381). The fraction of ‘narrowly’ tuned neurons – i.e., single-whisker unidirectional neurons – remained unchanged from L4 (0.081 ± 0.029, mean ± S.D., n=7 mice) to L3 (0.081 ± 0.016; L4 vs. L3, p = 0.957, paired t-test, n=7 mice) but declined in L2 (0.048 ± 0.022; **Fig. 3B**; L3 vs. L2, p < 0.001). Next, we examined the distribution of neuron types in layer-normalized depth (**Fig. 3C**) finding that unidirectional neuron frequency peaks in L4, bidirectional in L3, and multi-whisker in L2. Finally, we examined changes in encoding score (**Fig. 3D**). Unidirectional neurons showed consistently low encoding scores. In contrast, bidirectional neurons and multi-whisker neurons showed large increases in encoding score, implying that responses became more reliably predictable from whisker curvature.

**Figure 3.**
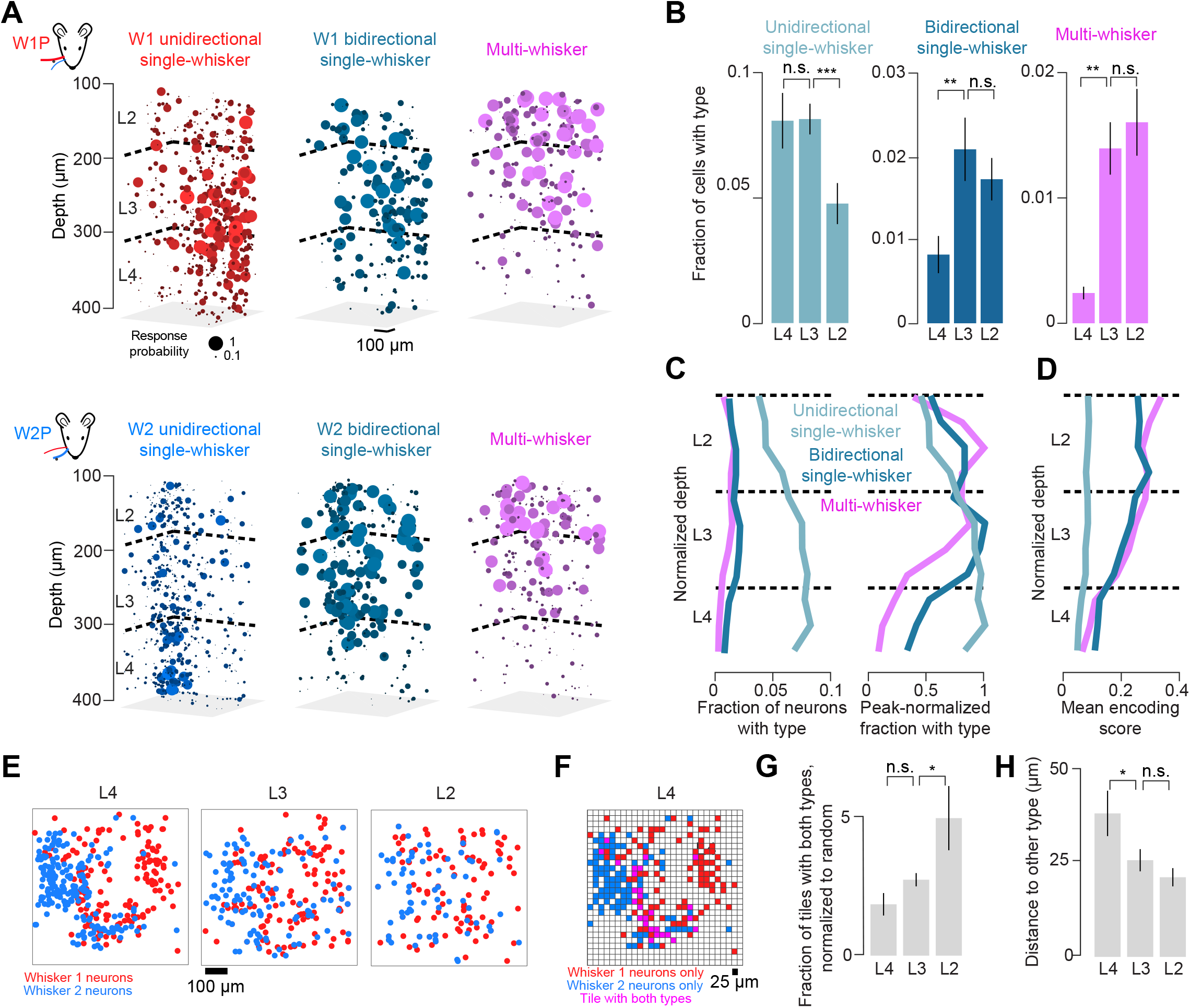
Frequency of multi-whisker neurons increases superficially as representations become interdigitated. **A)** Example map from one mouse showing the probability of response for W1P (top) and W2P (bottom) touch. Left to right: unidirectional single-whisker neurons, bidirectional single-whisker neurons, and multi-whisker neurons. **B)** Frequency of each of the major touch neuron types for L4, L3, and L2. Bars, mean; lines, S.E.M. (n=7). P-values indicated for paired t-test, *: p < 0.05; **: p < 0.01; ***: p < 0.001. **C)** Distribution of neuron types as a function of normalized depth (Methods). Left, fraction of neurons at a given depth. Right, within-type normalized fraction. **D)** Encoding score as a function of normalized depth. **E)** Distribution of single-whisker touch neurons in L2, L3, and L4 for an example mouse, projected onto the tangential plane. Red, whisker 1 neurons; blue, whisker 2 neurons. Multi-whisker and non-encoding neurons are omitted. **F)** Example tiling (25-by-25 μm) for the single-whisker L4 neurons from **E**. Tiles are colored based on the touch neurons they contain (red, whisker 1; blue, whisker 2; purple, both). **G)** Fraction of tiles having both neuron types across all layers and mice, normalized to the product of the single-whisker tile fractions. Bars, mean; lines, S.E.M. (n=7 mice). P-values as in **B**. **H)** Minimal distance for each layer between single-whisker neurons preferring different whiskers.

We next compared the distribution of unidirectional single-whisker neurons tuned to different touch directions – protraction and retraction - across layers. The fraction of retraction-preferring unidirectional neurons exceeded protraction preferring neurons in all layers, though the difference was not significant in L2 (**Fig S6A**). The decline from L4 to L2 for both protraction and retraction preferring neurons followed a similar pattern and encoding scores for both were similar across depths (**Fig. S6B, C**).

Connection probability in vS1 L2/3 increases with proximity between pairs of neurons (Holmgren et al 2003, Lefort et al 2009), so a superficial increase in the frequency of broadly tuned neurons could be due to more intermingling among narrowly tuned neurons. L3, and especially L2, exhibited greater intermingling of the two single-whisker representations than L4 (**Fig. 3E**). To quantify this, we projected all neurons from a layer onto a single plane that we then partitioned into 25-by-25 μm tiles. Tiles containing neurons responding to only one whisker were scored as belonging to that whisker, whereas tiles containing neurons responding to both whiskers were assigned to the multi-whisker group (**Fig. 3F**). We found that the ratio of multi-whisker tiles to the random prediction obtained by multiplying the frequencies of single-whisker tiles increased from L3 to L2 but not L4 to L3, (ratio, L4: 1.9 ± 1.1, L3: 2.8 ± 0.6, L2: 5.0 ± 3.1; L4 vs. L3, p = 0.156, paired t-test, n=7 mice; L3 vs. L2, p = 0.047; **Fig. 3G**; Methods). Further, the distance between a neuron responding to one whisker and the closest neuron responding to the other whisker decreased from L4 to L3 but not L3 to L2 (L4: 38 ± 16 μm, L3: 26 ± 8 μm, L2: 21 ± 6 μm; L4 vs. L3, p = 0.017; L3 vs. L2, p = 0.113; **Fig. 3H**). Thus, the superficial emergence of broadly tuned neurons is accompanied by greater spatial intermingling of narrowly tuned neurons.

### Superficial population response is sparser but more reliable

The increase in encoding score for multi-whisker and bidirectional neurons from L4 to L2 (**Fig. 3D**) suggests that superficial neurons are more consistently driven by touch. We therefore asked whether the probability of response to touch as well as the proportion of neurons that reliably respond to touch changed from L4 to L2. Focusing on the two most numerous strong (top third of Δκ) single-whisker touch types, W1P and W2P (**Fig. 1F, I**), we compared both the size of the responsive pool, defined as neurons with a touch response probability exceeding 0.1, and the response probability of neurons in this pool across layers. L4 responses to single-whisker touches were mostly confined to the barrel of the touching whisker, with many neurons exhibiting low response probability (**Fig. 4A**). L2 contained fewer responsive neurons, but they exhibited higher response probability across trials and were spatially dispersed. L3 exhibited an intermediate pattern. The fraction of neurons in the responsive pool was 0.16 ± 0.05 in L4 (mean ± S.D., n=7 mice) and 0.14 ± 0.05 in L3 (L4 vs. L3, p = 0.725, paired t-test, n=7 mice), dropping to 0.09 ± 0.05 in L2 (**Fig. 4B**; L3 vs. L2, p = 0.003). In contrast, the probability of response among response pool members increased from 0.13 ± 0.03 in L4 to 0.21 ± 0.04 in L3 (**Fig. 4B**; L4 vs. L3, p < 0.001), remaining unchanged at 0.22 ± 0.03 in L2 (L3 vs. L2, p = 0.308). Thus, from L4 to L2, the subset of neurons that responded declined, but the probability of response for any responsive neuron increased.

**Figure 4.**
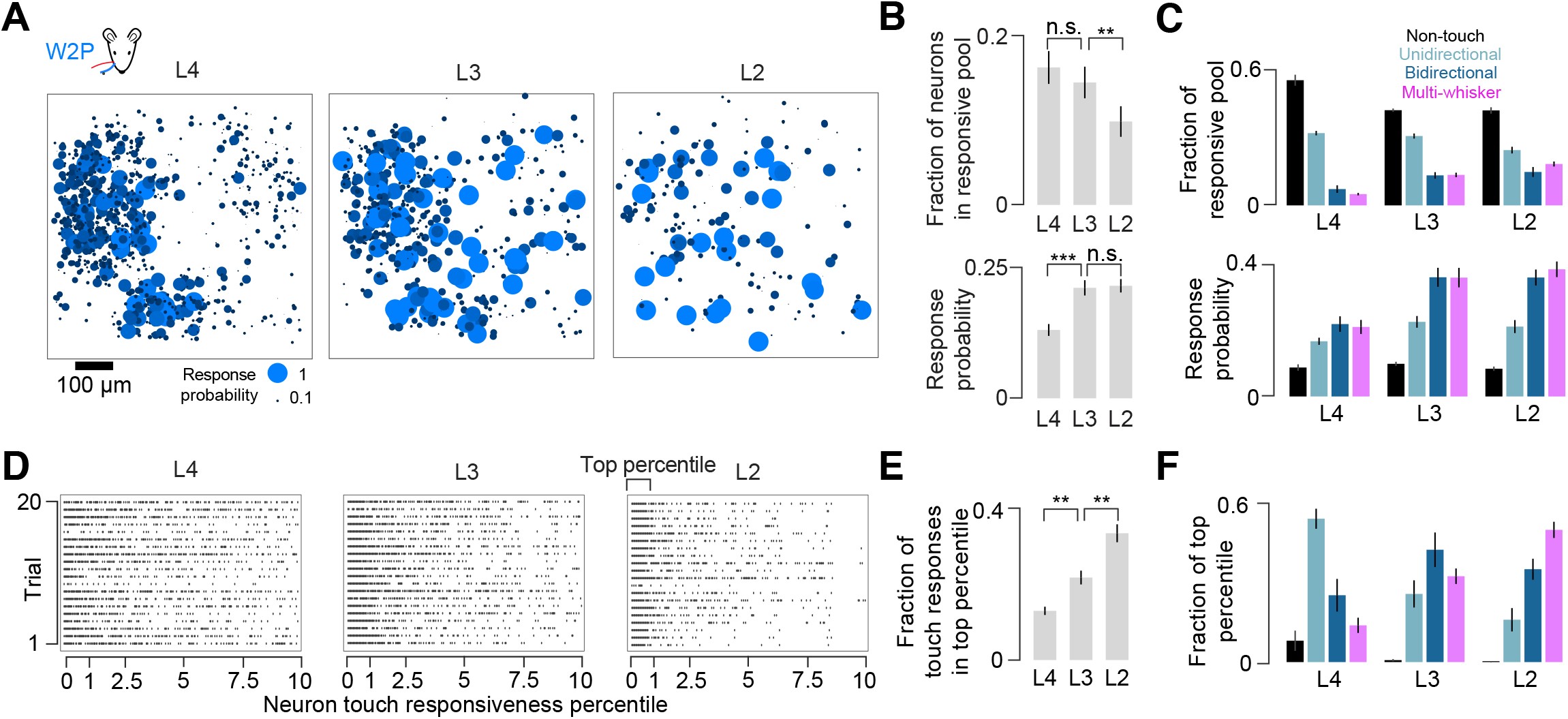
Transition to sparser, more reliable responses from L4 to L2. **A)** Population response to strong W2P touches across layers. Ball size and color indicate response probability (Methods). All neurons for a layer are projected onto the tangential plane. **B)** Response characteristics across layers. Top, fraction of neurons responding to strong W1P or W2P touches across layers (‘responsive pool’; Methods). Bottom, response probability across layers for neurons in this responsive pool. **C)** Responsiveness among different neural types. Top, fractional composition of the responsive pool in each layer. Bottom, probability of response for a given neuron type and layer. **D)** Example W1P responses for 20 strong touch trials, one trial per row, one group of 20 trials per layer. Neurons are sorted left-to-right by responsiveness percentile. A dot indicates that a particular neuron responded on a given trial. **E)** Fraction of touch-evoked calcium activity that originates from the top percentile of touch-responsive neurons. **F)** Neuron types comprising the most responsive percentile of neurons.

We next analyzed the composition of the responsive pool across layers. In L4, non-touch and unidirectional single-whisker (uSW) neurons made up a larger fraction of the responsive pool than bidirectional single-whisker (bSW) and multi-whisker (MW) neurons (**Fig. 4C**; p < 0.001, post-hoc Tukey’s HSD comparing any pair of neuron types within a layer, n=7 mice, except MW vs. bSW, p = 0.751). This pattern was maintained in L3 (p < 0.001 for all pairs except MW vs. bSW, p = 0.999), weakening slightly in L2 (p < 0.001 for all pairs except MW vs. bSW, p = 0.415, uSW vs. bSW, p = 0.001, and uSW vs. MW, p = 0.053). Nevertheless, the relative fraction of broadly tuned neurons increased from L4 to L2 while non-touch and narrowly tuned touch neurons came to make up a smaller portion of the response pool. Specifically, the fraction of the responsive pool consisting of broadly tuned neurons increased from L4 to L3 (bSW L4 vs. L3, p = 0.046, paired t-test, n=7 mice; MW, p < 0.001), with multi-whisker neurons further increasing in fraction from L3 to L2 (MW L4 vs. L3, p = 0.002; bSW, p = 0.549). The fraction of the responsive pool consisting of non-touch neurons declined from L4 to L3 (non-touch L4 vs. L3, p < 0.001; uSW, p = 0.491), and the fraction consisting of unidirectional single-whisker neurons declined from L3 to L2 (uSW L3 vs. L2, p = 0.003; non-touch, p = 0.665). Thus, broadly tuned neurons come to make up an ever-larger fraction of the responsive pool from L4 to L2.

Response probability also changed from L4 to L2 in a touch-class specific manner. In L4, response probability for touch neurons exceeded that for non-touch neurons (**Fig. 4C**; non-touch vs. uSW, p = 0.017, post-hoc Tukey’s HSD comparing neuron type pairs within a layer, n=7 mice; non-touch vs. bSW, p < 0.001; non-touch vs. MW, p < 0.001), whereas responses among different touch neuron classes were similar (uSW vs. bSW, p = 0.177; uSW vs. MW, p = 0.314; bSw vs. MW, p = 0.984). In contrast, in L3, broadly tuned neurons became more responsive than both non-touch neurons and unidirectional single-whisker neurons (non-touch vs. bSW, p < 0.001; non-touch vs. MW, p < 0.001; uSW vs. bSW, p = 0.001; uSW vs. MW, p = 0.002). This relationship was preserved in L2 (non-touch vs. bSW, p < 0.001; non-touch vs. MW, p < 0.001; uSW vs. bSW, p < 0.001; uSW vs. MW, p < 0.001). Thus, the responsiveness of broadly tuned neurons increases superficially, so that their activity becomes disproportionately impactful during touch.

To determine if the response was becoming more concentrated in these broadly tuned neurons, we examined the fraction of touch-evoked calcium events in the most touch responsive percentile of neurons (**Fig. 4D**; Methods). These neurons produce an increasing fraction of touch evoked calcium events from L4 to L3, and L3 to L2 (**Fig. 4E**; L4, fraction of response in top percentile: 0.13 ± 0.03, L3: 0.21 ± 0.05, L2: 0.33 ± 0.06; L4 vs. L3, p = 0.009, paired t-test, n=7 mice; L3 vs. L2, p = 0.001). This concentration of neural response is accompanied by a shift in composition among the top percentile, with a decline in the fraction of unidirectional single-whisker neurons and an increase in the fraction of multi-whisker neurons (**Fig. 4F**; uSW, L4 vs. L3, p = 0.001, paired t-test, n=7 mice; L3 vs. L2, p = 0.294; MW, L4 vs. L3, p = 0.001; L3 vs. L2, p = 0.001).

Thus, the transition to a sparser representation from L4 to L2 is accompanied by the emergence of a small group of broadly tuned neurons that respond more consistently to touch.

### Decoding for higher-order stimulus features improves from L4 to L2

Is this superficial increase in response reliability accompanied by improved stimulus decoding? To address this, we examined the decoding of touch features across layers.

We first examined the decoding of touch force. Specifically, for a given single whisker touch type, we asked how well neurons distinguish the strongest touches from the weakest touches using receiver operating characteristic (ROC) analysis (Methods; **Fig. 5A**). For a given single-whisker touch type, multi-whisker, bidirectional single-whisker, and unidirectional single-whisker neurons performed comparably in all layers (**Fig. 5B, C**; L4, p = 0.737, ANOVA comparing three neuron types within a layer, n=7 mice; L3, p = 0.317; L2, p = 0.708). We also did not observe differences in decoding of force for a single whisker from L4 to L3 or L3 to L2 across any touch neuron type (**Fig. 5C**). In contrast, multi-whisker and bidirectional single-whisker neurons outperformed unidirectional single-whisker touch neurons in most layers when decoding ability was assessed across all four single-whisker touch types (**Fig. 5D**; L4: MW vs. uSW, p < 0.001, post-hoc Tukey’s HSD comparing neuron types within a layer, n=7 mice; bSW vs. uSW, p = 0.010; L3: MW vs. uSW, p < 0.001; bSW vs. uSW, p = 0.028; L2: MW vs. uSW, p < 0.001; bSW vs. uSW, p = 0.113). Further, while decoding of strong vs. weak touches only improved marginally for unidirectional single-whisker neurons from L4 to L3 (uSW, p = 0.003, paired t-test, n=7 mice) and not at all from L3 to L2 (p = 0.749), it improved substantially from L4 to L3 for bidirectional and multi-whisker neurons (bSW L4 vs. L3, p = 0.004; L3 vs. L2, p = 0.095; MW L4 vs. L3, p = 0.009; L3 vs. L2, p = 0.099).

**Figure 5.**
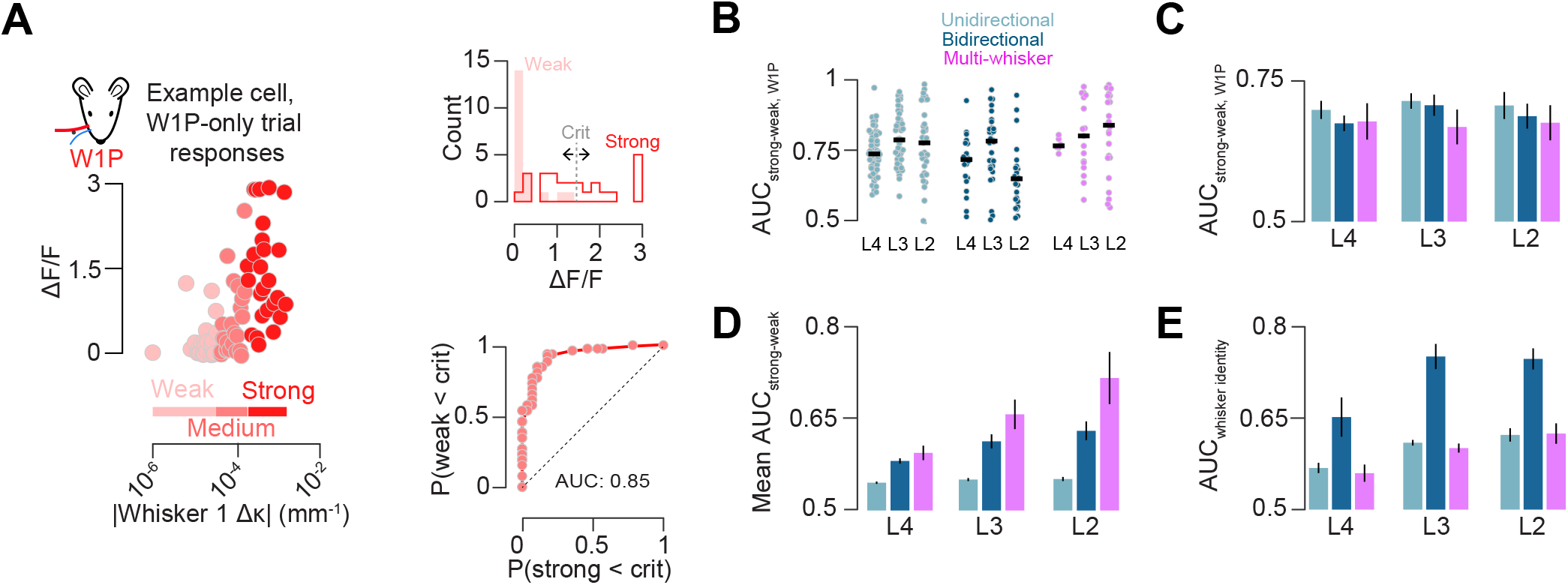
Decoding of touch features by neuron type and layer. **A)** Example ROC analysis for single neuron touch force decoding (Methods). Left, peak ΔF/F responses to W1P (red), as a function of Δκ. Protraction responses are divided into weak, medium, or strong thirds based on Δκ. Top right, histogram of strong and weak ΔF/F responses, with example criterion point for generating ROC curve. Bottom right, example neuron ROC curve. **B)** Decoding for all neurons from an example mouse of a given type and layer comparing neural response to strong and weak W1P touches. Dark line, median. **C)** AUC across all mice for W1P touch force. Bar indicates cross-animal mean, line indicates S.E.M. (n=7). **D)** As in **C**, but pooled across all four single whisker trial types. **E)** Discriminability of whisker identity.

We next asked how well individual neurons can distinguish between the two touching whiskers. We found that bidirectional neurons outperformed unidirectional and multi-whisker neurons across most layers (**Fig. 5E**; L4: bSW vs. MW p = 0.027, post-hoc Tukey’s HSD, n=7 mice; bSW vs. uSW, p = 0.066; L3: bSW vs. MW p < 0.001; bSW vs. uSW, p < 0.001; L2: bSW vs. MW, p < 0.001; bSW vs. uSW, p < 0.001). Among bidirectional single-whisker neurons, decoding improved from L4 to L3 (p = 0.010, paired t-test, bSW, n=7 mice) but not L3 to L2 (p = 0.818).

In sum, touch neurons of any type and layer perform comparably when decoding the force of force for a specific single whisker. Multi-whisker neurons perform best for decoding force in a whisker-invariant manner whereas bidirectional single-whisker neurons are best at decoding the identity of the touching whisker, with both exhibiting improved decoding from L4 to L3. Thus, the transition from L4 to L2 is accompanied by the emergence of specific functional processing streams with improved decoding of higher-order stimulus features and stable decoding of simple stimulus features.

We next asked how well populations of a given touch neuron type can decode touch strength for individual touch types (**Fig. S7A**). Decoding of weak vs. strong touch improved with population size, and broadly tuned populations performed as well as narrowly tuned populations (**Fig. S7B**). We asked how well groups of 10 neurons could decode touch across touch neuron types and layers. In contrast to single neurons, populations of broadly tuned neurons did not significantly outperform narrowly tuned neurons in decoding contact force across multiple touch types (**Fig. S7C**; L4 uSW vs. bSW, p = 0.583, paired t-test, n=7 mice; L3, p = 0.588, ANOVA comparing three touch neuron types, n=7 mice; L2, p = 0.196). Though the advantage of bidirectional neurons in discriminating whisker 1 touches from whisker 2 touches was smaller for the 10 neuron case than for single neurons, they performed better than other neural types in L3 and L2 (**Fig. S7D**; L3: bSW vs. MW, p < 0.001, post-hoc Tukey’s HSD, n=7 mice; bSW vs. uSW, p = 0.012; L2: bSW vs. MW, p = 0.031; bSW vs. uSW, p = 0.028), but not L4 (**Fig. S7D**; L4 uSW vs. bSW, p = 0.064, paired t-test, n=7 mice).

Thus, groups of unidirectional single-whisker neurons can achieve force decoding comparable to individual multi-whisker neurons and whisker identity decoding comparable to individual bidirectional neurons. Elevated decoding ability among individual broadly tuned neurons in L2/3 may therefore be due to pooling across multiple unidirectional single whisker neurons.

### Superficial broadly tuned neurons exhibit elevated functional coupling

Local axonal branches and dendritic arbors of rodent vS1 pyramidal neurons become more horizontally extensive from deep L3 to superficial L2 (Staiger et al 2014). If these reflect connectivity, we should observe coupling across a greater spatial extent superficially. We first examined pairwise correlations across all time points for neurons at various depths in our dataset. In L3 and L2, many bidirectional single-whisker neuron and multi-whisker neuron pairs exhibited high correlations (**Fig. 6A**). Broadly tuned neurons had higher pairwise correlations than single-whisker unidirectional touch neurons in L4 (**Fig. 6B**; bSW vs. uSW, p < 0.001, paired t-test, n=7 mice), L3 (bSW vs. uSW, p = 0.003, Tukey’s HSD, n=7 mice; MW vs. uSW, p = 0.016), and L2 (bSW vs. uSW, p = 0.001; MW vs. uSW, p = 0.001). In L3 and L2, bidirectional single-whisker neurons and multi-whisker neurons had comparable pairwise correlations (L3: MW vs. bSW, p = 0.753; L2: MW vs. bSW, p = 0.918). Correlations among bidirectional single-whisker neurons increased from L4 to L3 (L4 vs. L3, p = 0.003, paired t-test, n=7 mice), as did correlations among unidirectional single-whisker neurons (L4 vs. L3, p = 0.002), but neither changed from L3 to L2 (uSW, L3 vs. L2, p = 0.290; bSW, L3 vs. L2, p = 0.226). They increased slightly for multi-whisker neurons from L3 to L2 (L3 vs. L2, p = 0.025). Thus, pairwise correlations were highest for superficial broadly tuned neurons.

**Figure 6.**
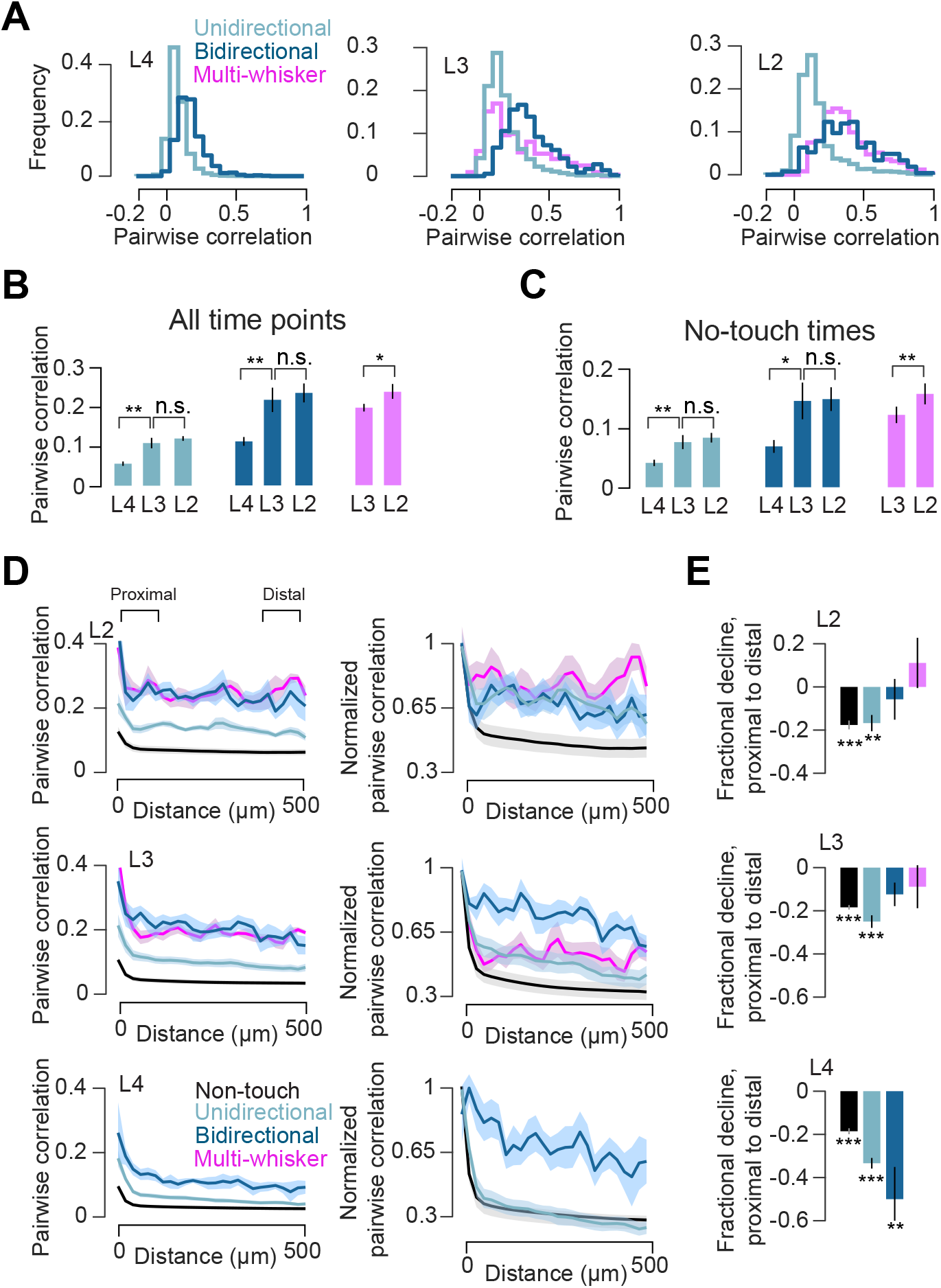
Pairwise correlations across layers and neuron types. **A)** Distribution of pairwise correlations for different touch types for an example animal. **B)** Average pairwise correlations for specific neuronal types, layers. Correlations were computed using all available imaging time. Bars, mean; lines, S.E.M. (n=7 mice). P-values indicated for paired t-test, *: p < 0.05; **: p < 0.01; ***: p < 0.001. **C)** As in **B**, but epochs around touch (from 1s prior to 10 s after) were excluded. **D)** Left, pairwise correlation as a function of distance. Right, within-animal normalized pairwise correlation as a function of distance. Shading indicates S.E.M. **E)** Fractional change in pairwise correlation from proximal to distal pairs. P-values indicated for paired t-test, proximal vs. distal, *: p < 0.05; **: p < 0.01; ***: p < 0.001.

Spontaneous activity often resembles stimulus-evoked activity (Kenet et al 2003, Luczak et al 2009) which modeling work shows may be due to underlying connectivity (Litwin-Kumar & Doiron 2014). We therefore examined the correlation structure during periods of no touch by excluding all time points from 1 s before to 10 s after any touch. Though the correlations were lower than those observed when including touch (**Fig. 6B**), the trends were mostly consistent. Unidirectional single-whisker neurons exhibited the lowest correlations in L4 (**Fig. 6C**; bSW vs. uSW, p = 0.011, paired t-test, n=7 mice) and L2 (bSW vs. uSW, p = 0.027, Tukey’s HSD, n=7 mice; MW vs. uSW, p = 0.012), and multi-whisker and bidirectional single-whisker neurons did not differ significantly (L2: MW vs. bSW, p = 0.921). Unlike correlations using all time, we did not find any differences among touch neuron types in L3 (p = 0.080, ANOVA). As with correlations measured for all time, non-touch correlations increased from L4 to L3 (uSW, L4 vs. L3, p = 0.005, paired t-test, n=7 mice; bSW, L4 vs. L3, p = 0.013), remained unchanged from L3 to L2 for single-whisker neurons (uSW, L3 vs. L2, p = 0.357; bSW, L3 vs. L2, p = 0.908), and increased slightly for multi-whisker neurons (L3 vs. L2, p = 0.010).

In sum, while correlations for all touch neuron types increase from L4 to L3, only multi-whisker correlations change from L3 to L2, increasing slightly. Broadly tuned neurons exhibit higher pairwise correlations than narrowly tuned neurons across layers.

We next asked if neurons exhibited elevated correlations across larger distances superficially, as predicted from morphology (Staiger et al 2014). Across all touch neuron types and layers, there is an initial decline in pairwise correlations over the first ~50 μm (**Fig. 6D**). As this represents a tiny fraction of all pairs, we compared correlations for pairs up to 100 μm apart (‘proximal’) to those 400-500 μm apart (‘distal’). Among non-touch neurons, the decline in pairwise correlations from proximal to distal was significant in all layers (**Fig. 6E**; L4, L3, L2: proximal vs. distal correlation, p < 0.001, paired t-test, n=7 mice). Unidirectional single-whisker neurons also showed a significant decline in correlation from proximal to distal in all layers (L4, L3: proximal vs. distal correlation, p < 0.001; L2, p = 0.007). Bidirectional single-whisker neurons, in contrast, only showed significant correlation decline with distance in L4 (L4: p = 0.004; L3: p = 0.072; L2: p = 0.570). Similarly, multi-whisker neurons did not show significant distance-dependent declines in correlation in L3 or L2 (L3: proximal vs. distal correlation, p = 0.408; L2: p = 0.384).

In sum, from L4 to L2, the distance-dependent drop in pairwise correlations among touch neurons weakens, with broadly tuned touch neurons exhibiting no drop in L3 or L2. Only non-touch neurons show a consistent distance-dependent drop in pairwise correlations across layers. Thus, functional coupling becomes more spatially extensive superficially for touch neurons, but not non-touch neurons.

### Superficial touch activity is concentrated in sparse groups of co-active touch neurons

Ensembles – groups of neurons that are co-active, often in response to a common stimulus – are frequently observed in cortex (Carrillo-Reid et al 2017, Harris & Mrsic-Flogel 2013). Given the elevated correlations among broadly tuned neurons, we asked if broadly tuned neurons are more likely to participate in ensembles. We assigned neurons to ensembles based on the pairwise correlation matrix across all simultaneously imaged pairs, grouping neurons whose mutual pairwise correlations were above a threshold (Methods; **Fig. 7A**). Some ensembles consisted almost entirely of touch neurons, with specific ensembles preferring specific types of touch (**Fig. 7B, C**); ensembles where 50% or more of neurons were touch neurons were considered ‘touch ensembles’.

**Figure 7.**
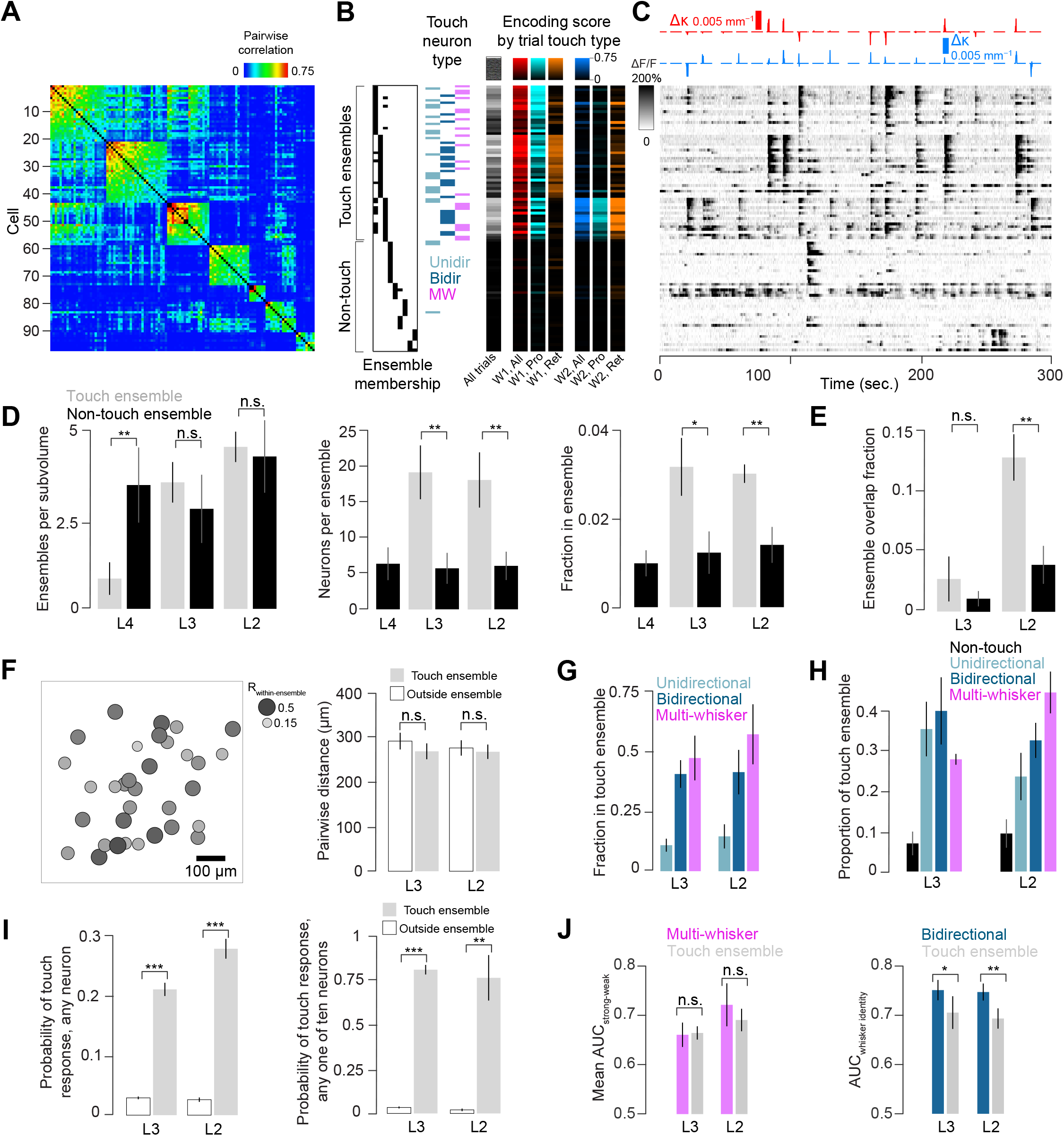
Ensembles and their role in touch responses. **A)** Pairwise correlation matrix for neurons in L2 of an example mouse that belong to ensembles (Methods). Ensembles are sorted by mean within-ensemble correlation. **B)** Classification of ensemble members. Left, assignment of each neuron to a particular ensemble. Here, the first three ensembles are touch ensembles. Middle, neural type. Right, encoding scores for given trial types. **C)** ΔF/F for the neurons in **A, B** over the course of 5 minutes. Top, Δκ for whisker 1 (red) and whisker 2 (blue). **D)** Basic properties of ensembles across layers. Left to right: ensembles per subvolume; number of neurons per ensemble; fraction of neurons belonging to an ensemble. Grey, touch ensembles. Black, non-touch ensembles. Bars indicate mean ± S.E.M. (n=7 mice). P-values indicated for paired t-test, *: p < 0.05; **: p < 0.01; ***: p < 0.001. **E)** Fractional overlap between ensembles. **F)** Left, spatial layout of the first touch ensemble in **A**. Right, mean pairwise distance for neurons within (grey) or outside (white) of touch ensembles (mean ± S.E.M., n=7 mice). **G)** Fraction of neurons belonging to given touch category that are also part of a touch ensemble. **H)** Composition of touch ensembles. **I)** Robustness of touch ensemble response. Left, probability of responding on any touch for a neuron within (grey) or outside (white) touch ensembles. Right, probability that at least one of ten randomly selected neurons from within or outside touch ensembles will respond to touch. **J)** Decoding by touch ensembles. Left, mean AUC for decoding strong vs. weak touches. Right, AUC for decoding touching whisker identity.

The number of touch ensembles per subvolume increased from 0.9 ± 1.2 (mean ± S.D., n=7 mice) in L4 to 3.6 ± 1.5 in L3 (L4 vs. L3, p = 0.004, paired t-test, n=7 mice), reaching 4.6 ± 1.1 in L2 (**Fig. 7D**; L3 vs. L2, p = 0.086). In contrast, the number of non-touch ensembles did not change, reaching 3.7 ± 3.0 in L4, 3.0 ± 2.7 in L3 (L4 vs. L3, p = 0.489), and 4.6 ± 2.9 in L2 (L3 vs. L2, p = 0.130). Non-touch ensembles were more numerous in L4 (touch vs. non-touch, p = 0.020, paired t-test, n=7 mice), but counts were comparable in L3 (p = 0.596) and L2 (p = 1). Because 4/7 mice had no touch ensembles in L4, we excluded L4 touch ensembles from subsequent analyses. We next asked whether ensemble size changed across layers. Touch ensembles in L3 contained 19.3 ± 10.0 neurons, similar to L2 touch ensembles, which contained 18.3 ± 10.3 neurons (L3 vs. L2, p = 0.844). Non-touch ensembles also remained stable in size, with 6.4 ± 6.1 neurons in L4, 5.8 ± 5.8 neurons in L3 (L4 vs. L3, p = 0.808), and 6.1 ± 5.3 neurons in L2 (L3 vs. L2, p = 0.874). Non-touch ensembles were smaller than touch ensembles in L3 (non-touch vs. touch ensemble neuron count, p = 0.003, n=7 mice) and L2 (p = 0.003). As with neuron count, the fraction of neurons in touch ensembles did not change from L3 (0.032 ± 0.018) to L2 (0.030 ± 0.006; L3 vs. L2, paired t-test, p = 0.800). Non-touch ensemble fraction also did not change from L4 (0.010 ± 0.008) to L3 (0.012 ± 0.013; L4 vs. L3, p = 0.608) or L3 to L2 (0.014 ± 0.011; L3 vs. L2, p = 0.686). The fraction of neurons in touch ensembles exceeded the non-touch ensemble fraction in L3 (non-touch vs. touch ensemble fraction, p = 0.014, n=7 mice) and L2 (p = 0.005). Thus, touch ensembles are mostly a feature of L3 and L2, with little difference in ensemble size or count between these layers; non-touch ensembles appear in all layers, are substantially smaller than touch ensembles, and exhibit little change between layers.

Individual neurons can belong to multiple ensembles (**Fig. 7B**). We quantified ensemble overlap as the size of the intersection among two ensembles divided by the size of their union. Overlap among touch ensembles increased from 0.03 ± 0.05 in L3 to 0.13 ± 0.05 in L2 (**Fig. 7E**; L3 vs. L2, p = 0.007, paired t-test, n=7 mice); the change in overlap among non-touch ensembles was not significant (L3, overlap: 0.01 ± 0.02; L2: 0.04 ± 0.04, L3 vs. L2, p = 0.133). Touch ensembles exhibited more overlap than non-touch ensembles in L2 (touch vs. non-touch, p = 0.001), but not L3 (p = 0.249). Thus, touch ensemble overlap increases from L3 to L2, and only exceeds non-touch ensemble overlap in L2.

Do touch ensemble neurons exhibit spatial clustering? To address this, we compared the mean pairwise distance between touch ensemble members to the distance between outside-ensemble pairs (**Fig. 7F**). We found no difference in L3 (ensemble vs. outside, 271 ± 47 vs. 294 ± 50 μm, p = 0.054, paired t-test, n=7 mice) or L2 (270 ± 43 vs. 278 ± 46 μm, p = 0.177). Thus, as with touch neurons (Peron et al 2015b), touch ensemble neurons do not exhibit spatial clustering.

Given the elevated pairwise correlations among broadly tuned neurons (**Fig. 6B, C**), we next asked whether these neurons participated in ensembles to a disproportionate degree. Throughout L2/3, broadly tuned neurons were indeed the most likely to participate in touch ensembles (**Fig. 7G**). Unidirectional single-whisker neurons were least likely to be part of touch ensembles (L3 MW vs. uSW, p = 0.002, Tukey’s HSD, n=7 mice; bSW vs. uSW, p = 0.016; L2 MW vs. uSW, p = 0.009; bSW vs. uSW, p = 0.017). We did not observe any change from L3 to L2 for the proportion of various touch neuron types belonging to ensembles (L3 vs. L2, MW, p = 0.565, paired t-test, n=7 mice; bSW, p = 0.974; uSW, p = 0.861). Thus, despite their rarity, nearly half of broadly tuned neurons in L2/3 participate in ensembles.

Though the fraction of a given touch neuron type participating in touch ensembles did not change from L3 to L2, the change in composition between these two layers (**Fig. 3B**) suggests that broadly tuned neurons should come to make up a larger portion of ensembles in L2. We therefore looked at touch neuron types that made up ensembles across layers, finding that the fraction of ensemble neurons that were also multi-whisker neurons increased from L3 to L2 (**Fig. 7H**; L3 vs. L2, p-value = 0.032, paired t-test, n=7 mice), while the unidirectional single-whisker neuron fraction declined (**Fig. 7H**; L3 vs. L2, p-value = 0.018). Bidirectional single-whisker fraction did not change (L3 vs. L2, p-value = 0.366), nor did the non-touch fraction (L3 vs. L2, p-value = 0.428). Thus, though the three touch neuron types contribute comparably to touch ensembles in L3, in L2, multi-whisker neurons come to dominate at the expense of unidirectional single-whisker neurons.

We next asked how reliably touch ensemble neurons responded to touch. Touch ensemble neurons were more reliable in responding to touch than neurons outside touch ensembles, especially superficially (**Fig. 7I**). Specifically, the probability of any touch ensemble neurons responding to touch exceeded that of neurons outside touch ensembles for L3 (touch ensemble response probability: 0.21 ± 0.03, outside ensemble: 0.03 ± 0.01, ensemble vs. outside ensemble, p < 0.001, paired t-test, n=7 mice) and L2 (touch ensemble response probability: 0.28 ± 0.04, outside ensemble: 0.03 ± 0.01, p < 0.001). The response probability for touch ensembles increased from L3 to L2 (p-value = 0.009). With a larger pool of neurons, the difference between touch ensemble and outside ensemble neurons increased even more: the probability that at least one neuron of 10 responds to a touch exceeds the outside-ensemble probability in L3 (within ensemble: 0.81 ± 0.07, outside ensemble: 0.03 ± 0.01, p < 0.001, paired t-test, n=7 mice) and L2 (within ensemble: 0.77 ± 0.34, outside ensemble: 0.02 ± 0.02, p = 0.001). Thus, downstream readout pooling across a small number (~10) of touch ensemble members could reliably detect the majority of touches. Given this reliability, we asked how well touch ensemble neurons decode touch features. We found that touch ensemble neurons decoded strong from weak touches across all touch types as well as multi-whisker neurons (**Fig. 7J**; L3, touch ensemble vs. multi-whisker decoding, p = 0.871 paired t-test, n=7 mice; L2, p = 0.329). When decoding touch whisker identity, however, bidirectional single-whisker neurons slightly but significantly outperformed touch ensemble neurons (**Fig. 7J**; L3, touch ensemble vs. bidirectional single-whisker decoding, p = 0.031; L2: p = 0.009). Thus, only certain stimulus features are more effectively represented by ensembles.

## DISCUSSION

We assess trans-laminar transformations in cortical layers 2-4 of vS1 by densely sampling neural activity in mice performing a two whiskers object localization task. We observe distinct depth distributions for different functional neuron types: unidirectional single-whisker neurons peak in L4, bidirectional single-whisker neurons in L3, and multi-whisker neurons in L2. This is accompanied by a transition from touch responses in L4 in which individual neurons respond more randomly to touch but the population of potential responders is large, to sparser but more consistent responsiveness in L3 and especially L2. Decoding of force across multiple whiskers as well as decoding of whisker identity improve from L4 to L2, with specific touch neuron types best decoding each feature; decoding of force from a single whisker remains high across all layers. Superficially, we find ensembles (Carrillo-Reid et al 2017, Harris & Mrsic-Flogel 2013) of broadly tuned touch-responsive neurons that, despite making up a small minority of the population (~ 3 % of neurons) constitute a large and highly reliable portion of the responsive population (> 25 %) on any given touch. Thus, the transition from L4 to L2 yields sparser yet more informative and robust responses, potentially facilitating perceptual readout (Barlow 1972, Sakurai 1999, Wolfe et al 2010).

Layers 2 and 3 are often treated as a single layer. Though many differences between L2 and L3 exist (Petersen & Crochet 2013), physiological and morphological changes are often gradual (Staiger et al 2014). Consequently, even the depth of the L2-L3 boundary within rodent vS1 has been debated, with some authors proposing a thinner L2 and thicker L3 (Hooks et al 2011, Narayanan et al 2017), whereas others define them to be of approximately equal size (Lefort et al 2009). Dividing L2/3 equally, we find several differences between L2 and L3. First, the fraction of unidirectional single-whisker neurons declines from L3 to L2. Second, spatial intermingling among neurons tuned to different whiskers is more pronounced in L2 than L3. Third, the fraction of neurons responding to touch declines, and the fraction of touch-evoked activity occurring in the most responsive neurons increases, so that L2 touch responses are more concentrated than L3. Thus, changes in connectivity and morphology from deep L3 to superficial L2 (Staiger et al 2014) are accompanied by specific changes in sensory responses.

Changes in receptive field structure have been observed from L4 to L2 in many primary sensory cortices, typically in anesthetized preparations (Kanold et al 2014, Martinez et al 2005, Simons 1978). In V1, there is an increase in the number of complex cells relative to simple cells in L2/3 (Hubel & Wiesel 1962, Martinez et al 2005, Ringach et al 2002, Van Hooser et al 2013), and in A1, L2/3 neurons have higher frequency bandwidth in their tuning compared to L4 neurons (Winkowski & Kanold 2013). Our observed broadening of vibrissal receptive fields in superficial vS1 demonstrates that such broadening is not just a consequence of anesthesia, which expands receptive field size in the vibrissal system (Friedberg et al 1999, Simons et al 1992). We find two forms of broadening superficially: the emergence of bidirectional neurons, and the emergence of multi-whisker neurons. In contrast, single-whisker unidirectional neurons decline in frequency superficially.

What is the circuit basis for receptive field broadening in superficial vS1? Spines in L2 vS1 neurons exhibit a mix of single- and multi-whisker responses (Varga et al 2011), but the relative contributions of intra-laminar input, input from L4 (Lefort et al 2009), or even direct thalamic input (Audette et al 2017) remain unknown. Given the distance-dependence of connections (Holmgren et al 2003) and the abundance of intra-laminar connections (Lefort et al 2009), the observed spatial broadening in connectivity from L4 to L2 (Staiger et al 2014) likely contributes to the decline in single-whisker unidirectional neurons from L4 to L2 and increasing frequency of more broadly tuned neurons. At the same time, multi-whisker input from posterior medial thalamus preferentially targeting L2 (Jouhanneau et al 2014), and even feedback from higher order areas (Lee et al 2013, Nandy et al 2017), also likely play a role. Future experiments will be needed to elucidate the contributions of specific inputs to L2/3 receptive fields.

Our decoding analysis suggests the two broadly tuned populations perform specific functions. Multi-whisker neurons can accurately decode contact force across multiple whiskers, with improved decoding from L4 to L2. This emergence of a whisker invariant force code suggests that for these neurons, the vibrissal system optimizes the extraction of contact force at the expense of whisker identity. Whisker identity, however, is well-represented by the bidirectional single-whisker neurons, with decoding improving from L4 to L2. In contrast, decoding of single whisker touch force is already high in L4 and does not improve superficially. Thus, from L4 to L2, decoding of features poorly decoded by L4 improves, while preserving accurate decoding of features which L4 decodes well.

We remove all but two of our animals’ whiskers. Many neurons we classify as single whisker are therefore likely to have multi-whisker receptive fields. Thus, while our overall touch neuron fraction (10 %) is consistent with previous estimates (Crochet et al 2011, O’Connor et al 2010b, Peron et al 2015b), our multi-whisker neuron fraction is lower (Clancy et al 2015). The transition from unidirectional and bidirectional single-whisker neurons to multi-whisker neurons is likely to be even more pronounced than reported here.

We did not observe neurons that showed responsiveness exclusively to multi-whisker contacts, despite observations of enhanced responses when whiskers are stimulated in rapid succession under other contexts (Laboy-Juarez et al 2019). Such responses are likely crucial in texture processing, where slip-stick events occur as the animal’s whisker moves against a surface (Jadhav & Feldman 2010, Jadhav et al 2009). Because our task employs active touch, the animal dictates the inter-touch interval. Thus, in contrast to studies employing direct stimulation of multiple whiskers (Laboy-Juarez et al 2019), we observed relatively few inter-touch intervals below 50 ms, the typical range where enhancement is seen.

Sparse activity is a common feature of sensory L2/3, yet its origin and function remain unclear (Barth & Poulet 2012, Harris & Mrsic-Flogel 2013, Wolfe et al 2010). We find that sparse populations of touch neurons in L2/3 of vS1 exhibit high pairwise correlations, suggesting that the responding neurons are interconnected, as they are in L2/3 of mouse V1 (Cossell et al 2015, Lee et al 2016, Wertz et al 2015). Several lines of evidence support this view. First, sparse populations in vS1 expressing the activity-linked immediate early gene c-*fos* exhibit elevated connectivity (Yassin et al 2010). Second, touch evoked responses in L2/3 (Crochet et al 2011) fall within the time window of maximal synaptic potentiation (Banerjee et al 2014), so that repeated touch should drive connectivity among these neurons. Finally, following the lesion of tens of touch neurons, the spared touch population shows a decline in responsiveness consistent with recurrent amplification (Peron et al 2020). Given that synchronous excitatory activity evokes strong feedback inhibition in vS1 L2/3 (Chettih & Harvey 2019, Dalgleish et al 2020, Kapfer et al 2007, Mateo et al 2011), sparseness may be a consequence of co-active groups of neurons that suppress activity among the remaining neurons via feedback inhibition (Barth & Poulet 2012, Wolfe et al 2010). Our observation of touch ensembles containing neurons with high pairwise correlations and robust touch responses suggests that sparseness is a consequence of the coactivation of such groups of neurons. Simultaneously, we find that touch ensemble members show both robust decoding of vibrissal features and highly reliable touch responses. This suggests that sparsification may be a key function of superficial cortical circuitry, yielding a small population of neurons that provides a robust stimulus response well-suited for perceptual readout (Barlow 1972, Wolfe et al 2010).

We show that from L4 to L2, the touch population response transitions from a diffuse and probabilistic one consisting mostly of narrowly tuned neurons to a sparse and robust one consisting mostly of broadly tuned neurons organized into ensembles. In L2/3 of mouse V1, stimulation of a small number of ensemble neurons can drive perceptual report, suggesting that small groups of such neurons can strongly influence perception (Carrillo-Reid et al 2019, Marshel et al 2019). Though we do not explicitly test the perceptual role of touch ensemble neurons, our decoding analysis suggests that feedforward processing in superficial cortex improves decoding for certain stimulus features, thereby facilitating their perceptual readout (Barlow 1972, Wolfe et al 2010). The sparse nature of vS1 ensemble responses makes them ideal candidates for cellular-resolution perturbation experiments (Emiliani et al 2015) testing their perceptual role and exploring the circuit basis of their responses.

## ACKNOWLEDGMENTS

We thank Adam Carter, Michael Long, and Robert Shapley for comments on the manuscript. We thank Ravi Pancholi and Lauren Ryan for discussion. This work was supported by the Whitehall Foundation and the National Institutes of Health (R01NS117536; R90DA043849).

## AUTHOR CONTRIBUTIONS

B.V. and S.P. designed the study. B.V. carried out all the experiments. B.V. and S.P. performed data analysis and wrote the paper.

## METHODS

### RESOURCE AVAILABILITY

#### Data and Code Availability

Source code used in this paper will be made available at http://github.com/peronlab following publication. Data from this paper will be made available in a public repository following publication. Requests for either prior to publication should be directed to.

### EXPERIMENTAL MODEL AND SUBJECT DETAILS

#### Animals and Surgery

Cranial windows were assembled by gluing a 3.5 mm circular #1.5 coverslip to a 4.5 mm circular #1.5 coverslip (Norland 61 glue). Windows were implanted over vS1 in P60-P90 Ai162 (JAX 031562) X Slc17a7-Cre (JAX X 023527) mice (Daigle et al 2018) of mixed sex, as described previously (Peron et al 2020). In vS1, these mice express GCaMP6s exclusively in excitatory neurons. Following surgical recovery, mice were placed on water restriction. The location in vS1 of barrels corresponding to whiskers C1-3 was identified by measuring the ΔF/F at coarse resolution (4X; 2.2 x 2.2 mm field of view) on a two-photon microscope while the whiskers were individually deflected. Animals were trimmed to the two whiskers whose barrels had the least obstructive vasculature, typically C2 and C3. Subsequent trimming occurred every 2-3 days. All animal procedures were in compliance with protocols approved by New York University’s University Animal Welfare Committee.

#### Behavior

Following surgical recovery, mice were water restricted. Next, mice were handled and head-fixed so as to habituate to the behavioral apparatus. Mice were trained on an object localization task (Peron et al 2020) in which a metal pole (0.5 mm diameter; Drummund Scientific, PA, USA) enters into the range of the mouse’s whiskers either at a distal position or at a range of proximal positions, typically spanning 5 mm along the anterior-posterior axis. On any given proximal trial, the pole appears at random position drawn from the range. In all trial types, the pole remains within the whisking plane for 2 s, after which it is moved back out of reach, below the whisking plane. 0.5 s after the pole is withdrawn, an auditory cue (3.4 kHz, 50 ms) indicates to the mouse to make a response, with the left lickport rewarded on distal trials and the right lickport rewarded on proximal trials. On all trials, the lickport is withdrawn and moves into an accessible position only during the response epoch (i.e., after the auditory cue). Incorrect responses result in a timeout and premature withdrawal of the lickport. Mice were considered to reach criterion performance once d-prime exceeded 1.5 for two consecutive days.

#### Whisker videography

Whisker video was acquired using custom MATLAB (MathWorks) software from a CMOS camera (Ace-Python 500, Basler) running at 400 Hz and 640 x 352 pixels and using a telecentric lens (TitanTL, Edmund Optics). Illumination was via a pulsed 940 nm LED (SL162, Advanced Illumination). 7-8s of each trial were imaged, including 1s prior to pole movement, the period when the pole was in reach, and several seconds after the pole was retracted. Data was processed on NYU’s High Performance Computing (HPC) cluster: first, candidate whiskers were detected using the Janelia Whisker Tracker (Clack et al 2012). Next, whisker identity was refined and assessed across a single session using custom MATLAB software (Peron et al 2020, Peron et al 2015b). Following whisker assignment, curvature (κ) and angle (θ) were calculated at specific locations along each whisker’s length. Change in curvature, Δκ, was calculated relative a resting angle-dependent baseline curvature value obtained during periods when the pole was out of reach. Next, automatic touch detection was performed. Touch assignment was manually curated using a custom MATLAB user interface (Peron et al 2020).

With the exception of our touch-detection algorithm, analyses, including model fitting, employed a down-sampled version of Δκ to match the sampling rate of the calcium imaging data (7 Hz vs. 400 Hz). Specifically, we used the maximal |Δκ| value over the ~140 ms duration of a single imaging frame, while preserving the sign. To obtain a single Δκ value for a trial, we computed the mean across all time points for that trial during which the whisker is touching the pole. Where applicable, trials were partitioned into equal sized thirds based on this value, resulting in ‘strong’, ‘medium’, and ‘weak’ touch trial groupings. For multi-whisker touch trials, the trial was assigned a type on the basis of the first two distinct single-whisker touches in that trial (**Fig. 1F, I**).

#### Two-photon imaging

Imaging was performed using a MIMMS (http://openwiki.janelia.org/wiki/display/shareddesigns/MIMMS) two-photon microscope with a 16X objective (Nikon). Illumination was at 940 nm (Chameleon Ultra 2; Coherent), with power rarely exceeding 50 mW. Three imaging planes spanning 700-by-700 μm (512-by-512 pixels) and spaced 20 μm apart (‘subvolume’) were acquired at a rate of ~7Hz. Depth was modulated with a piezo (P-725KHDS; Physik Instrumente). Power was depth-adjusted in software with an exponential length constant typically having a value of 250 μm. Imaging data was acquired using Scanimage (Vidrio Technologies).

Each of 5-7 subvolumes was imaged for 50-70 trials, followed by the next subvolume, and so on. Most subvolumes were imaged on any given day. After the first imaging day, motion-corrected mean images were collected for each plane and used as reference images on subsequent days.

Imaging data were processed on the NYU HPC cluster immediately after acquisition, as described previously (Peron et al 2015b). The first step was motion correction via image registration. Next, for the first day of imaging, neurons were detected using an automated algorithm based on template convolution. This initial segmentation was manually curated, and a reference segmentation was established for that plane. On subsequent imaging sessions, the reference segmentation was algorithmically transferred to the new data (Huber et al 2012). Following segmentation, neuropil subtraction and ΔF/F computation were performed. Finally, event detection was performed. For most analyses, the ΔF/F trace was used; calcium events were only used where explicitly mentioned.

#### Layer assignment

For each animal, a reference image with an interplane spacing of 2 μm was collected under light anesthesia (Isoflurane, ~1% by volume; **Fig. S1**). Stacks were started just above the dura. Because the dura provides a reliably strong elevated fluorescent signal, we could automatically detect its appearance. The stack was divided into a 5-by-5 grid in the imaging plane, and dura depth was determined for each segment of the grid. The resulting points were fit to a plane using the singular value decomposition and used as the surface of the brain. For each point in the reference stack, depth was assigned based on the distance along a line through the point and perpendicular to the dural surface plane. This corrected for the fact that the objective’s image plane was typically tilted with respect to the dural surface by several degrees.

As described above, we used a single reference plane, typically from the first day of imaging, to align our imaging during subsequent sessions. We fit this reference image to the stack volume using a thin plate spline fitting algorithm. This allowed us to obtain a common (x, y, depth) coordinate for each pixel in the plane and, by extension, assign a depth to each recorded neuron.

The L1-L2 border was defined as the depth of the most superficially imaged excitatory neuron. The L3-L4 border was found by manually locating a noticeable shift in neuron morphology in conjunction with the emergence of clearly visible septa. The L2-L3 border was placed at the midpoint between the L1-L2 and L3-L4 borders. L5 was not imaged in most mice; when it was, the L4-L5 border was defined as the first appearance of large somata, and all neurons at or beyond this depth were excluded from analysis. Because laminar boundaries are not discrete, analyses comparing layers were performed while excluding neurons within a 50 μm slice centered on the laminar boundary. This was done for the L2-L3 border and the L3-L4 border. Layers were typically thinner than observed anatomically (Lefort et al 2009), likely due to compression from the cranial window.

Normalized depth was obtained by assigning a depth of 0, 1, and 2 to the L1-L2, L2-L3, and L3-L4 borders, respectively. Since L2 and L3 were of equal thickness, this was just a simple rescaling from the depth of the L1-L2 and L3-L4 borders to 0 and 2. Neurons in L4 were assigned a normalized depth using: 2+(distance to L3-L4 border)/(distance from L2-L3 border to L3-L4 border).

#### Resolution of overlapping signal sources

Source contamination due to pixels containing multiple neurons is a concern in two-photon calcium imaging (Song et al 2021). This is especially problematic in the axial direction when a neuron appears across multiple planes. To identify such duplicates and to ensure that our data did not include multiple instances of the same neuron, we manually inspected all instances of candidate neuron pairs within a cylinder of radius 20 μm and half-height 40 μm that had a correlation above 0.2, keeping the neuron with the strongest signal and removing the others in instances where there were multiple candidates per real neuron. This resulted in the removal of 2,835 ± 962 candidate neurons per mouse (mean ± S.D.; n=7 mice).

#### Encoding model and neural classification

Neurons were classified based on how well an encoding model could predict their activity on specific trial types (**Fig. S4**). The model predicts neural activity (ΔF/F), *r*_*model*_, from

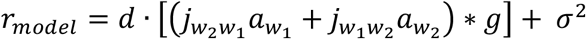

Where 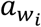 is the predicted amplitude of response to a given whisker at a given time, *g* is the GCaMP kinetics kernel for that neuron, *d* is a session-specific scaling factor, *j* is a cross-whisker interaction term, and *σ*^2^ is a Gaussian noise term. For single-whisker touch trials for whisker *i*, 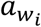 is

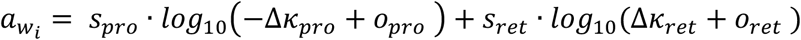

This model is based on previous work using a less constrained generalized linear model that revealed monotonically increasing response as a function of whisker curvature across touch neurons(Peron et al 2020). For a given whisker, the amplitude of the response to a protraction touch (*Δκ* < 0) at a given time, 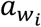, is given by applying a slope *s*_*pro*_ to its change in curvature, *Δκ*_*pro*_. To account for neurons that have a minimal force needed to elicit a response, the offset term *o*_*pro*_ was included. The retraction (*Δκ* > 0) response is calculated in an analogous manner.

The indicator kinetics kernel, *g*, consisted of a sum of exponentials having time constants *τ*_*rise*_ and *τ*_*decay*_. It was normalized so that its peak was 1. Both *τ*_*rise*_ and *τ*_*decay*_ were constrained based on the known physiological range (Chen et al 2013): *τ*_*rise*_, 100 ms to 500 ms; *τ*_*decay*_,1 s to 5 s. The noise term *σ*^*2*^ was determined for each neuron by measuring the variance of negative ΔF/F values. Our sliding-window F_0_ fitting procedure (Peron et al 2015b), in which we compute F_0_ using a 3 minute sliding window as the median for neurons that have low activity (non-skewed F distribution) and the 5^th^ percentile for the most active neurons (highly skewed F distribution) ensures that ΔF/F is appropriately 0-centered.

The linear scaling factor *d* assumed a single, unique value per session. Across all sessions, the maximal value of *d* was set to 1. For all other sessions, *d* reached a value between 0 and 1. This was designed to absorb variation in response across the multiple days of imaging.

The model was fit with 5-fold cross validation using block coordinate descent and a mean-square-error cost function minimizing the difference between model response, *r*_*model*_, and neural response, *r*_*neural*_. During cross-validation, data was partitioned by randomly drawing 5 disjoint equal sized sets of trials; individual trials were not broken up. The terms of 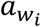 were iteratively fit along with *g* and *d* using single-whisker touch trials and an equal number of non-touch trials. Because this resulted in two estimates of *g* and *d*, we employed the mean of these parameters for the final model fit; manual inspection revealed that the two individual whisker fits predicted similar values for these terms.

Following the single-whisker fits, a second fitting procedure was run to fit the interaction terms 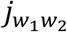 and 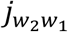 using the trials where both whiskers touched. Multi-whisker touch trials were classified based on the first two touches, and the scaling factor was applied to all touches by the second-touching whisker on that trial. Only trials where the interval between these first two touches was < 200 ms were included; this typically included the majority of multi-whisker trials (**Fig. S5A**). For any trial where whisker 1 touched first, 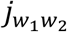 was allowed to vary from 0 to 10, with values below 1 corresponding to suppression of the whisker 2 response and values above 1 corresponding to enhancement. An analogous procedure was used for trials where whisker 2 touched first.

The output of the full model, *r*_*model*_, was a predicted ΔF/F trace. In addition, a shuffled fit was performed in which the ΔF/F was shifted temporally, with wrap-around, by a random number of timesteps (minimum: 10 s, the approximate length of a trial) while leaving the Δκ vectors untouched. A single shuffled fit was performed per neuron. To obtain a distribution of shuffled fits, neurons in a subvolume were grouped into 10 equal bins based on their calcium event rate, and neurons in a given bin used all shuffled fits in that bin. We classified neurons by using the Pearson correlation of the model-predicted and actual ΔF/F traces for specific trial types. Thus, a unidirectional single-whisker neuron would be one whose ΔF/F trace and model-predicted ΔF/F trace had a correlation for a single touch direction’s trials that met two criteria: correlation in excess of 0.1 and exceeding the 99^th^ percentile of shuffled data correlation for event rate matched neurons. Neurons were classified as unidirectional single-whisker neurons if they only met criteria when the Pearson correlation was calculated for one single-whisker touch type (W1P, W1R, W2P, or W2R). Neurons meeting criteria for both touch types for a single whisker were classified as bidirectional single-whisker. Neurons were classified as multi-whisker if they met our criteria for at least one direction for each whisker.

#### Response probability analysis

Neurons were classified as responsive or non-responsive for every touch trial by comparing the post-touch ΔF/F to the baseline ΔF/F. Baseline ΔF/F was calculated as the mean ΔF/F for the 6 frames (0.85 s) preceding the first touch on that trial. The post-touch ΔF/F was calculated as the mean ΔF/F for the period between the first touch and two frames after the final touch. For each neuron, we obtained a noise estimate by fitting all negative ΔF/F values to a half-normal distribution, yielding a noise term, σ. Neurons were considered responsive on a given trial if the ΔF/F_post-touch_ > ΔF/F_baseline_ + σ and if ΔF/F_post-touch_ exceeded the 99^th^ percentile of shuffled ΔF/F_post-touch_ values for that trial. Shuffled ΔF/F_post-touch_ was calculated by temporally shifting the ΔF/F vector, with wrap-around, by a random number of timesteps (at least 10 s, the approximate length of a trial). This was done 100 times per neuron, yielding a distribution of shuffled ΔF/F_post-touch_ values for each neuron and touch trial. Neurons that were responsive on at least 10 % of all touch trials for a given touch type were considered part of the response pool.

#### Decoding analysis

Single neuron decoding was performed by computing the post-touch ΔF/F across two sets of trials. We used two trial partitioning schemes. For force decoding, single-whisker trials of a single type (W1P, W1R, W2P, W2R) were divided into equal-sized thirds based on mean trial Δκ. The ability of a neuron to distinguish between the top third (‘strong’) and bottom third (‘weak’) of trials based on Δκ was evaluated. For whisker identity decoding, all single whisker touch trials for whisker 1 were compared to single whisker trials for whisker 2.

For each trial belonging to the pair of trial types under examination, we computed the mean ΔF/F between the first touch and first lick; if the first lick occurred > 2 s after the first touch, only 2 s after the first touch were used. Receiver operating characteristic (ROC) analysis was performed by sliding a criterion threshold through the range of ΔF/F values across the two trial types. We report the area under the curve (AUC) resulting from this analysis (Green & Swets 1966). Decoding was performed only if at least ten trials of each type were present.

Population decoding was performed by repeatedly drawing n_neurons_ from the specified population (1,000 repetitions). An estimate for confidence bounds for population decoding ability was obtained by randomizing trial labels for all neurons independently. In all cases, the mean ΔF/F value over the 2 s after touch was calculated for each neuron; concatenating these resulting in a population response vector of length n_neurons_ for a given trial. A decision variable was calculated, equal to the dot-product similarity to the mean population response vector for trials of the first type minus the dot-product similarity to the mean population response for trials of the second type (O’Connor et al 2010b). As with single neuron decoding, ROC analysis was performed using the distribution of this decision variable across the two trial types.

#### Correlation analysis

Pearson correlations were calculated across neuron pairs for each layer using the subvolume that had the most neurons belonging to that layer for that particular animal. We measured correlations either over all time points, or restricted to time points outside of touch. This was defined as excluding any timepoints 1 s prior to and 10 s after a touch. Pairwise distances were computed using the three-dimensional coordinates of the evaluated neurons. When computing correlations, single-whisker neurons were grouped by subtype. That is, for unidirectional single-whisker contacts, we independently computed a mean correlation for each animal (and, for distance analysis, for a given distance) among W1P, W1R, W2P, and W2R neurons. The mean of these was then used as the correlation for unidirectional single-whisker neurons. A similar process was used for bidirectional single-whisker neurons, with grouping by whisker. For multi-whisker neurons, we did not break up by subtype, as the number of neurons of a given subtype was too low. Consequently, correlation for multi-whisker neurons is likely underestimated. Despite this, we consistently find this group to exhibit the highest pairwise within-group correlations. Multi-whisker neurons were excluded from L4 because their number was small.

#### Ensemble detection

Ensembles were identified using a greedy algorithm. As with correlation and decoding analysis, the subvolume with the most neurons for a given layer was used in analyzing ensembles for that layer, allowing for analysis of concurrently recorded neurons. Hierarchical clustering was performed on the correlation matrix computed for all time points to identify small (3-5 neuron) groups of neurons with high mutual pairwise correlations. Each cluster was used as a seed for the greedy algorithm, which at each step added the neuron that had the highest mean correlation with the existing ensemble members. The process continued until no neuron could be added without reducing the mean within-ensemble correlation below a threshold defined as twice the 99.5^th^ percentile of all correlation values. Because this produced many redundant ensembles, ensembles with at least 75 % overlap were merged provided this would not violate the minimum correlation criteria. Ensembles with less than 3 members were discarded; such ensembles were rare, and never showed touch-related activity. The percent of touch neurons in an ensemble was most often either ~0 % or ~100 % (**Fig. 7B**); therefore, touch ensembles were defined as those ensembles for which at least 50 % of neurons were classified as touch neurons by our encoding model.

#### Statistical analysis

For comparisons across two matched groups, the paired t-test was used. Typically, pairing was within-animal. Multiple (3 or more) groups were compared using a one-way ANOVA. In cases where this yielded significance (p < 0.05), post-hoc Tukey’s Honestly Significant Difference (HSD) test is reported for comparisons between pairs of groups.

## REFERENCES

Adesnik H, Naka A. 2018. Cracking the Function of Layers in the Sensory Cortex. Neuron 100: 1028–43

Audette NJ, Urban-Ciecko J, Matsushita M, Barth AL. 2017. POm Thalamocortical Input Drives Layer-Specific Microcircuits in Somatosensory Cortex. Cereb Cortex: 1–17

Banerjee A, Gonzalez-Rueda A, Sampaio-Baptista C, Paulsen O, Rodriguez-Moreno A. 2014. Distinct mechanisms of spike timing-dependent LTD at vertical and horizontal inputs onto L2/3 pyramidal neurons in mouse barrel cortex. Physiological reports 2: e00271

Barlow HB. 1972. Single units and sensation: a neuron doctrine for perceptual psychology? Perception 1: 371–94

Barth AL, Poulet JF. 2012. Experimental evidence for sparse firing in the neocortex. Trends Neurosci 35: 345–55

Buzsaki G. 2010. Neural syntax: cell assemblies, synapsembles, and readers. Neuron 68: 362–85

Carrillo-Reid L, Han S, Yang W, Akrouh A, Yuste R. 2019. Controlling Visually Guided Behavior by Holographic Recalling of Cortical Ensembles. Cell 178: 447–57 e5

Carrillo-Reid L, Yang W, Kang Miller JE, Peterka DS, Yuste R. 2017. Imaging and Optically Manipulating Neuronal Ensembles. Annual review of biophysics 46: 271–93

Chen TW, Wardill TJ, Sun Y, Pulver SR, Renninger SL, et al. 2013. Ultrasensitive fluorescent proteins for imaging neuronal activity. Nature 499: 295–300

Chettih SN, Harvey CD. 2019. Single-neuron perturbations reveal feature-specific competition in V1. Nature 567: 334–40

Clack NG, O’Connor DH, Huber D, Petreanu L, Hires A, et al. 2012. Automated tracking of whiskers in videos of head fixed rodents. PLoS Comput Biol 8: e1002591

Clancy KB, Schnepel P, Rao AT, Feldman DE. 2015. Structure of a single whisker representation in layer 2 of mouse somatosensory cortex. J Neurosci 35: 3946–58

Constantinople CM, Bruno RM. 2013. Deep cortical layers are activated directly by thalamus. Science 340: 1591–4

Cossell L, Iacaruso MF, Muir DR, Houlton R, Sader EN, et al. 2015. Functional organization of excitatory synaptic strength in primary visual cortex. Nature 518: 399–403

Crochet S, Poulet JF, Kremer Y, Petersen CC. 2011. Synaptic mechanisms underlying sparse coding of active touch. Neuron 69: 1160–75

Daigle TL, Madisen L, Hage TA, Valley MT, Knoblich U, et al. 2018. A Suite of Transgenic Driver and Reporter Mouse Lines with Enhanced Brain-Cell-Type Targeting and Functionality. Cell 174: 465–80 e22

Dalgleish HW, Russell LE, Packer AM, Roth A, Gauld OM, et al. 2020. How many neurons are sufficient for perception of cortical activity? Elife 9

Douglas RJ, Martin KA. 2004. Neuronal circuits of the neocortex. Annu Rev Neurosci 27: 419–51

Emiliani V, Cohen AE, Deisseroth K, Hausser M. 2015. All-Optical Interrogation of Neural Circuits. J Neurosci 35: 13917–26

Friedberg MH, Lee SM, Ebner FF. 1999. Modulation of receptive field properties of thalamic somatosensory neurons by the depth of anesthesia. J Neurophysiol 81: 2243–52.

Green DM, Swets JA. 1966. Signal detection theory and psychophysics. New York: John Wiley & Sons Ltd.

Harris KD, Mrsic-Flogel TD. 2013. Cortical connectivity and sensory coding. Nature 503: 51–8

Hebb DO. 1949. The organization of behavior; a neuropsychological theory. New York: Wiley. xix, 335 p. pp.

Hirsch JA, Martinez LM. 2006. Laminar processing in the visual cortical column. Current opinion in neurobiology 16: 377–84

Holmgren C, Harkany T, Svennenfors B, Zilberter Y. 2003. Pyramidal cell communication within local networks in layer 2/3 of rat neocortex. J Physiol 551: 139–53

Hooks BM, Hires SA, Zhang YX, Huber D, Petreanu L, et al. 2011. Laminar analysis of excitatory local circuits in vibrissal motor and sensory cortical areas. PLoS Biol 9: e1000572

Hubel DH, Wiesel TN. 1962. Receptive fields, binocular interaction and functional architecture in the cat’s visual cortex. J. Physiol. (London) 160: 106–54

Huber D, Gutnisky DA, Peron S, O’Connor DH, Wiegert JS, et al. 2012. Multiple dynamic representations in the motor cortex during sensorimotor learning. Nature 484: 473–8

Jadhav SP, Feldman DE. 2010. Texture coding in the whisker system. Current opinion in neurobiology 20: 313–8

Jadhav SP, Wolfe J, Feldman DE. 2009. Sparse temporal coding of elementary tactile features during active whisker sensation. Nat Neurosci 12: 792–800

Ji XY, Zingg B, Mesik L, Xiao Z, Zhang LI, Tao HW. 2016. Thalamocortical Innervation Pattern in Mouse Auditory and Visual Cortex: Laminar and Cell-Type Specificity. Cereb Cortex 26: 2612–25

Jouhanneau JS, Ferrarese L, Estebanez L, Audette NJ, Brecht M, et al. 2014. Cortical fosGFP expression reveals broad receptive field excitatory neurons targeted by POm. Neuron 84: 1065–78

Kanold PO, Nelken I, Polley DB. 2014. Local versus global scales of organization in auditory cortex. Trends Neurosci 37: 502–10

Kapfer C, Glickfeld LL, Atallah BV, Scanziani M. 2007. Supralinear increase of recurrent inhibition during sparse activity in the somatosensory cortex. Nat Neurosci 10: 743–53

Kenet T, Bibitchkov D, Tsodyks M, Grinvald A, Arieli A. 2003. Spontaneously emerging cortical representations of visual attributes. Nature 425: 954–6

Kobatake E, Tanaka K. 1994. Neuronal selectivities to complex object features in the ventral visual pathway of the macaque cerebral cortex. J Neurophysiol 71: 856–67

Laboy-Juarez KJ, Langberg T, Ahn S, Feldman DE. 2019. Elementary motion sequence detectors in whisker somatosensory cortex. Nat Neurosci 22: 1438–49

Lee S, Kruglikov I, Huang ZJ, Fishell G, Rudy B. 2013. A disinhibitory circuit mediates motor integration in the somatosensory cortex. Nat Neurosci 16: 1662–70

Lee WC, Bonin V, Reed M, Graham BJ, Hood G, et al. 2016. Anatomy and function of an excitatory network in the visual cortex. Nature 532: 370–4

Lefort S, Tomm C, Floyd Sarria JC, Petersen CC. 2009. The excitatory neuronal network of the C2 barrel column in mouse primary somatosensory cortex. Neuron 61: 301–16

Litwin-Kumar A, Doiron B. 2014. Formation and maintenance of neuronal assemblies through synaptic plasticity. Nat Commun 5: 5319

Luczak A, Bartho P, Harris KD. 2009. Spontaneous events outline the realm of possible sensory responses in neocortical populations. Neuron 62: 413–25

Marshel JH, Kim YS, Machado TA, Quirin S, Benson B, et al. 2019. Cortical layer-specific critical dynamics triggering perception. Science 365

Martinez LM, Wang Q, Reid RC, Pillai C, Alonso JM, et al. 2005. Receptive field structure varies with layer in the primary visual cortex. Nat Neurosci 8: 372–9

Mateo C, Avermann M, Gentet LJ, Zhang F, Deisseroth K, Petersen CC. 2011. In vivo optogenetic stimulation of neocortical excitatory neurons drives brain-state-dependent inhibition. Curr Biol 21: 1593–602

Meyer HS, Wimmer VC, Hemberger M, Bruno RM, de Kock CP, et al. 2010. Cell type-specific thalamic innervation in a column of rat vibrissal cortex. Cereb Cortex 20: 2287–303

Mountcastle VB. 1997. The columnar organization of the neocortex. Brain 120 ( Pt 4): 701–22

Nandy AS, Nassi JJ, Reynolds JH. 2017. Laminar Organization of Attentional Modulation in Macaque Visual Area V4. Neuron 93: 235–46

Narayanan RT, Udvary D, Oberlaender M. 2017. Cell Type-Specific Structural Organization of the Six Layers in Rat Barrel Cortex. Frontiers in neuroanatomy 11: 91

Niell CM, Stryker MP. 2008. Highly selective receptive fields in mouse visual cortex. J Neurosci 28: 7520–36

O’Connor DH, Clack NG, Huber D, Komiyama T, Myers EW, Svoboda K. 2010a. Vibrissa-based object localization in head-fixed mice. J Neurosci 30: 1947–67

O’Connor DH, Peron SP, Huber D, Svoboda K. 2010b. Neural activity in barrel cortex underlying vibrissa-based object localization in mice. Neuron 67: 1048–61

Olshausen BA, Field DJ. 2004. Sparse coding of sensory inputs. Curr Opin Neurobiol 14: 481–7

Peron S, Chen TW, Svoboda K. 2015a. Comprehensive imaging of cortical networks. Curr Opin Neurobiol 32: 115–23

Peron S, Pancholi R, Voelcker B, Wittenbach JD, Olafsdottir HF, et al. 2020. Recurrent interactions in local cortical circuits. Nature 579: 256–59

Peron SP, Freeman J, Iyer V, Guo C, Svoboda K. 2015b. A Cellular Resolution Map of Barrel Cortex Activity during Tactile Behavior. Neuron 86: 783–99

Petersen CC, Crochet S. 2013. Synaptic computation and sensory processing in neocortical layer 2/3. Neuron 78: 28–48

Reddy L, Kanwisher N. 2006. Coding of visual objects in the ventral stream. Curr Opin Neurobiol 16: 408–14

Ringach DL, Shapley RM, Hawken MJ. 2002. Orientation selectivity in macaque V1: diversity and laminar dependence. J Neurosci 22: 5639–51

Sakata S, Harris KD. 2009. Laminar structure of spontaneous and sensory-evoked population activity in auditory cortex. Neuron 64: 404–18

Sakurai Y. 1999. How do cell assemblies encode information in the brain? Neurosci Biobehav Rev 23: 785–96

Severson KS, Xu D, Van de Loo M, Bai L, Ginty DD, O’Connor DH. 2017. Active Touch and Self-Motion Encoding by Merkel Cell-Associated Afferents. Neuron 94: 666–76 e9

Simons DJ. 1978. Response properties of vibrissa units in rat SI somatosensory neocortex. J Neurophysiol 41: 798–820

Simons DJ, Carvell GE, Hershey AE, Bryant DP. 1992. Responses of barrel cortex neurons in awake rats and effects of urethane anesthesia. Exp. Brain Res. 91: 259–72

Song A, Gauthier JL, Pillow JW, Tank DW, Charles AS. 2021. Neural anatomy and optical microscopy (NAOMi) simulation for evaluating calcium imaging methods. J Neurosci Methods 358: 109173

Staiger JF, Bojak I, Miceli S, Schubert D. 2014. A gradual depth-dependent change in connectivity features of supragranular pyramidal cells in rat barrel cortex. Brain structure & function

Van Hooser SD, Roy A, Rhodes HJ, Culp JH, Fitzpatrick D. 2013. Transformation of receptive field properties from lateral geniculate nucleus to superficial V1 in the tree shrew. J Neurosci 33: 11494–505

Varga Z, Jia H, Sakmann B, Konnerth A. 2011. Dendritic coding of multiple sensory inputs in single cortical neurons in vivo. Proceedings of the National Academy of Sciences of the United States of America 108: 15420–5

Wertz A, Trenholm S, Yonehara K, Hillier D, Raics Z, et al. 2015. PRESYNAPTIC NETWORKS. Single-cell-initiated monosynaptic tracing reveals layer-specific cortical network modules. Science 349: 70–4

Wimmer VC, Bruno RM, de Kock CP, Kuner T, Sakmann B. 2010. Dimensions of a projection column and architecture of VPM and POm axons in rat vibrissal cortex. Cereb Cortex 20: 2265–76

Winkowski DE, Kanold PO. 2013. Laminar transformation of frequency organization in auditory cortex. J Neurosci 33: 1498–508

Wolfe J, Houweling AR, Brecht M. 2010. Sparse and powerful cortical spikes. Curr Opin Neurobiol 20: 306–12

Woolsey TA, Van der Loos H. 1970. The structural organization of layer IV in the somatosensory region (SI) of mouse cerebral cortex. The description of a cortical field composed of discrete cytoarchitectonic units. Brain Res 17: 205–42

Yassin L, Benedetti BL, Jouhanneau JS, Wen JA, Poulet JF, Barth AL. 2010. An embedded subnetwork of highly active neurons in the neocortex. Neuron 68: 1043–50

