## Supplementary Material for "Transformation of Primary Sensory Cortical Representations from Layer 4 to Layer 2"

#### SUPPLEMENTARY MATERIALS

**Table S1. List of animals, Related to Figure 1.** All mice were transgenic Ai162 X Slc17a7-Cre (Daigle et al., 2018), expressing GCaMP6s in excitatory neurons. Age is given in days. Unlike for most analyses, where neurons in a 50 µm slice centered on the layer boundary were omitted, provided cell counts include all neurons imaged.

| Animal ID | Sex | Age of Training Start | Spared Whiskers | Number of Subvolumes | L2 Cell Count | L3 Cell Count | L4 Cell Count |
|---|---|---|---|---|---|---|---|
| **274688** | M | 67d | C1, C2 | 7 | 5,431 | 6,833 | 5,351 |
| **279029** | M | 73d | C2, C3 | 5 | 3,140 | 6,370 | 4,608 |
| **279608** | F | 59d | C2, C3 | 5 | 1,927 | 3,776 | 5,689 |
| **280201** | M | 81d | C2, C3 | 6 | 4,152 | 4,512 | 6,792 |
| **283544** | M | 93d | C2, C3 | 5 | 3,844 | 5,098 | 5,672 |
| **284891** | F | 60d | C2, C3 | 5 | 3,818 | 6,688 | 4,703 |
| **284893** | M | 63d | C1, C2 | 5 | 4,263 | 4,205 | 4,788 |

#### Voelcker and Peron, Figure S1

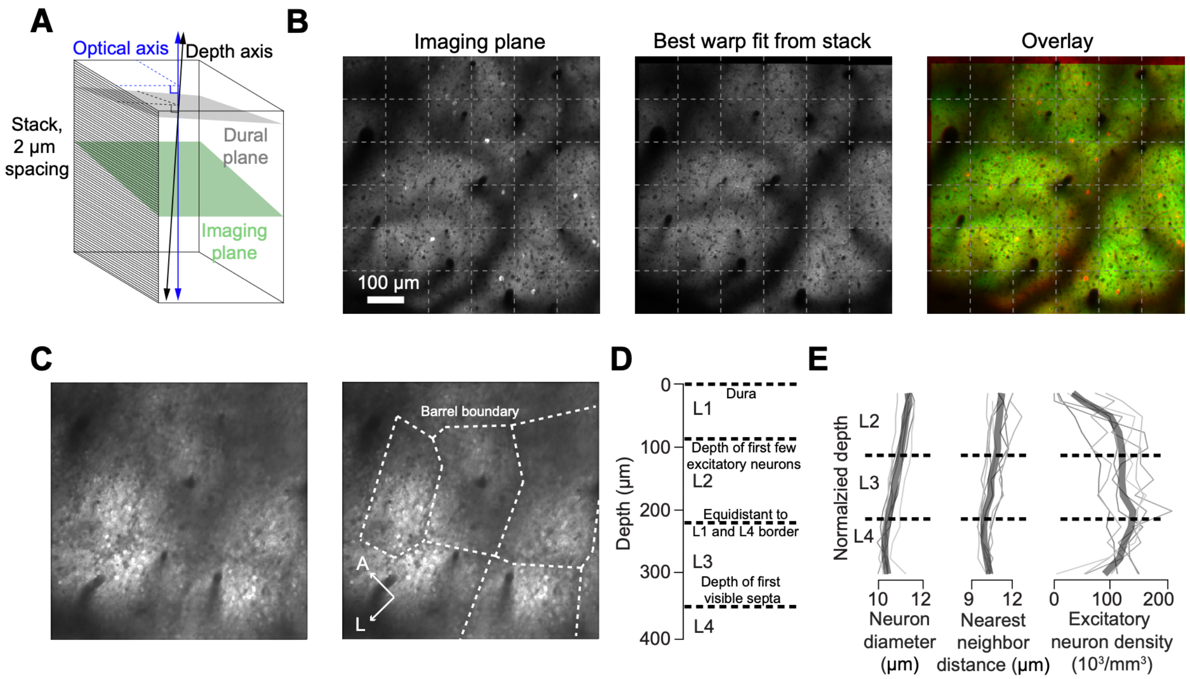

**Figure S1, Related to Figure 1. Assignment of cortical depth.**

**A)** A reference stack with 2 µm spacing is used to assign cortical depth. Depth is calculated with respect to the axis perpendicular to the dural plane (grey; Methods). The imaging plane (green) is then aligned to the reference stack, allowing for an assignment of depth to each imaging plane pixel.

**B)** Example thin plate spline warp field fit. Left, the imaging plane. Middle, best fit obtained from the stack. Due to anesthesia, activity is reduced in the stack plane. Right, overlay of both.

**C)** Example septa in L4; appearance of septa was used as criterion for determining L3-L4 border. Left, raw image. Right, barrel boundaries inferred from septa.

**D)** Method for the assignment of each laminar border.

**E)** Morphological parameters as a function of normalized depth. Light lines, individual animals; dark thick line, cross-animal mean. Left, mean diameter of cells at a given depth. Middle, distance for each neuron to its nearest neighbor. Right, volumetric density of excitatory neurons.

**Voelcker and Peron, Figure S2**

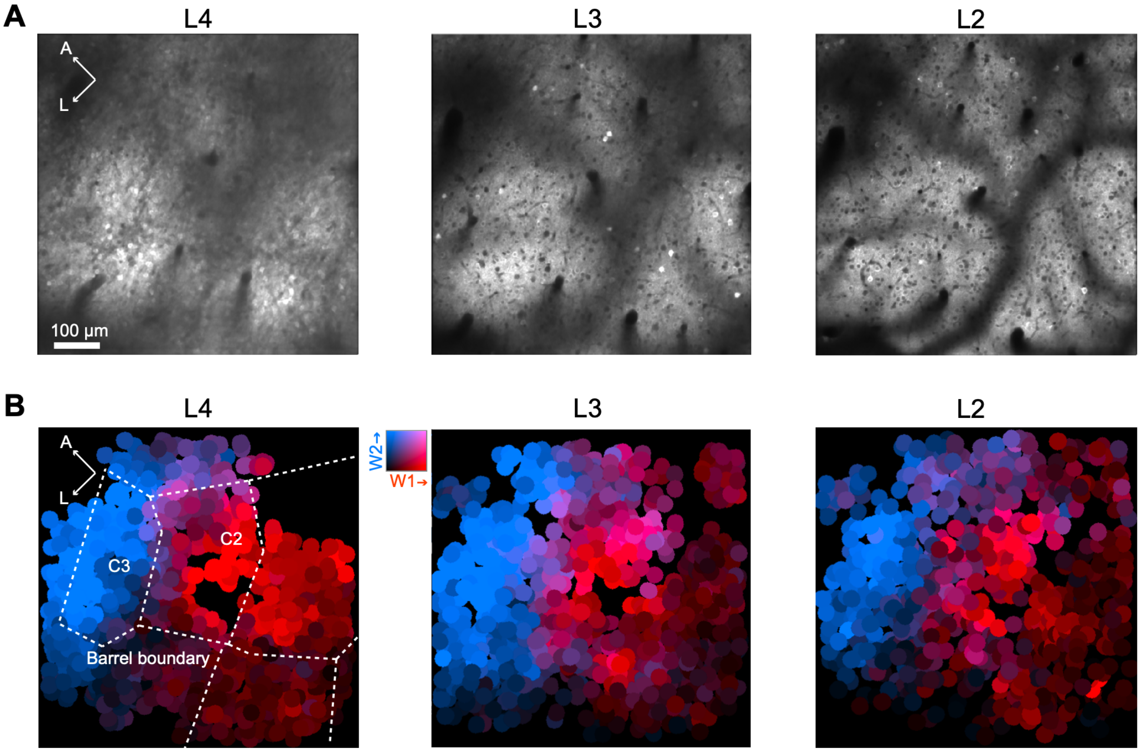

**Figure S2, Related to Figure 1. Barrel identification.**
**A)** Example planes from L2, L3 and L4.
**B)** Neuropil signal centered at each soma. As described previously, the neuropil signal was computed for all pixels within 3-13 µm away from the neuron border, excluding any pixels belonging to a soma or those with neighbor pairwise correlation exceeding 0.2 (Peron et al., 2015). Color code indicates sensitivity to whisker 1 (red) or whisker 2 (blue); in this case, the C2 and C3 whiskers. Barrel boundaries from L4 in **A** are traced in the whisker map.

Voelcker and Peron, Figure S3

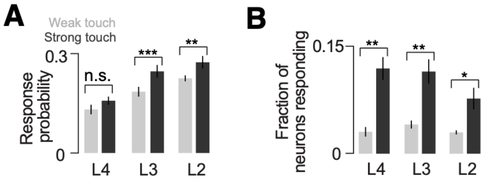

**Figure S3, Related to Figure 2. Population response increases with touch strength.**

**A)** Response probability for neurons in responsive pool across layers, touch intensities. P-values indicated for paired t-test, *: $p < 0.05$; **: $p < 0.01$; ***: $p < 0.001$.

**B)** Fraction of neurons that are part of the ‘responsive pool’ (i.e., neurons that respond on at least 10 % of touches) for a given touch strength.

**Voelcker and Peron, Figure S4**

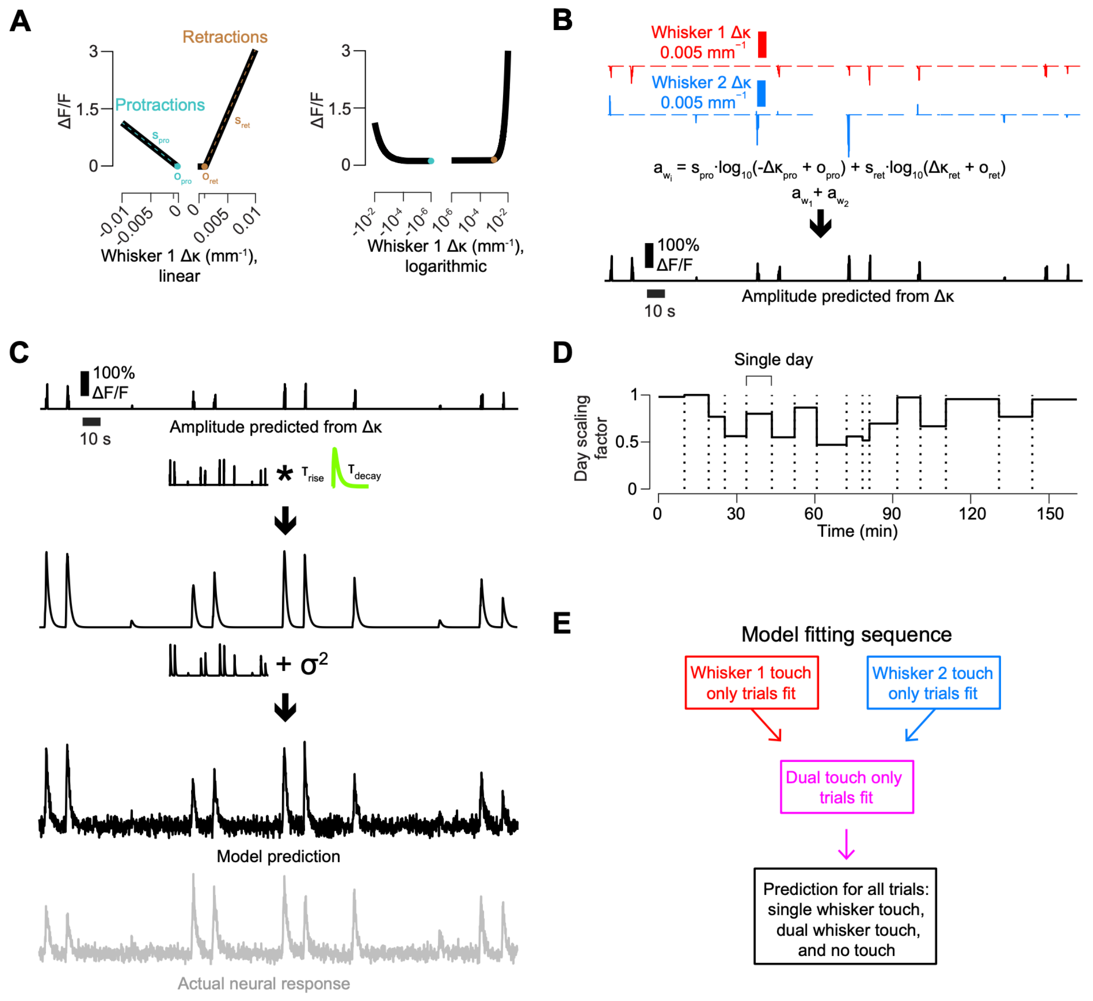

**Figure S4, Related to Figure 2. Encoding model.**

**A)** Example Δκ kernel, which maps instantaneous Δκ to a ΔF/F amplitude. If the cell has a Δκ threshold for that particular direction of touch, $o_{ret}$ or $o_{pro}$ will be non-zero. Right, example kernel in logarithmic coordinates.

**B)** The individual kernels are applied to the relevant Δκ trace to produce $a_{w_i}$ for each whisker; these are summed to produce the overall ΔF/F amplitude prediction.

**C)** The amplitude prediction from **B** is convolved with a GCaMP6s kinetics kernel, which is a sum of exponentials (Methods). Next, noise is added ($\sigma^2$; Methods), resulting in the full prediction.

**D)** Example day scaling factor applied over the imaging days.

**E)** Fitting procedure for exclusive touch model.

### Voelcker and Peron, Figure S5

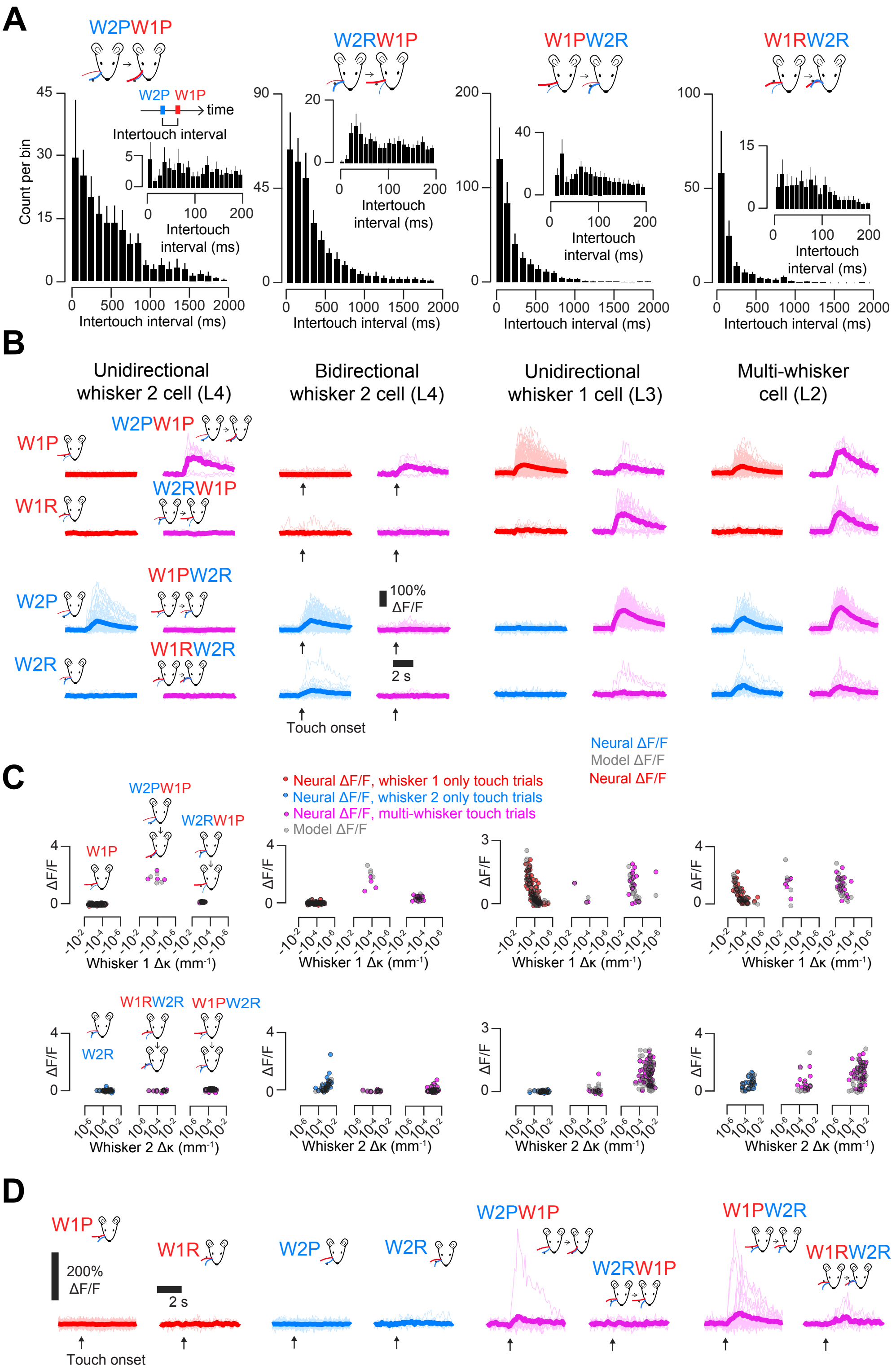

**Figure S5, Related to Figure 2. Responses on multi-whisker touch trials.**
**A)** Inter-touch interval distributions for the four multi-whisker touch types (W2PW1P, W2RW1P, W1PW2R, W1RW2R). The interval is between the first two touches on any multi-whisker trial; 100 ms bins were used, with inset showing 10 ms bins for first 200 ms. Bar shows mean ± S.E.M. (n=7 mice). Plot includes all touches for a given mouse.
**B)** Example ΔF/F responses to four single-whisker and four multi-whisker touch types for the four neurons from **Fig. 2A, E**. Light color, individual touch-aligned responses; dark color, mean across touches. Traces are colored according to touch type, indicated above the traces.
**C)** Comparison of actual vs. model tuning curves. The mean ΔF/F as a function of Δκ for each trial is shown with colored circles. Red circles, trials where only whisker 1 touched; blue, only whisker 2 touched; purple, both whiskers touched. Gray circles, model's predicted ΔF/F for the same trials. For multi-whisker touch trials, Δκ is given for the second whisker that touched. By convention, Δκ is negative for protraction touches.
**D)** One of two neurons across the entire dataset that showed no response to single whisker touches but substantial responses to multi-whisker touches.

Voelcker and Peron, Figure S6

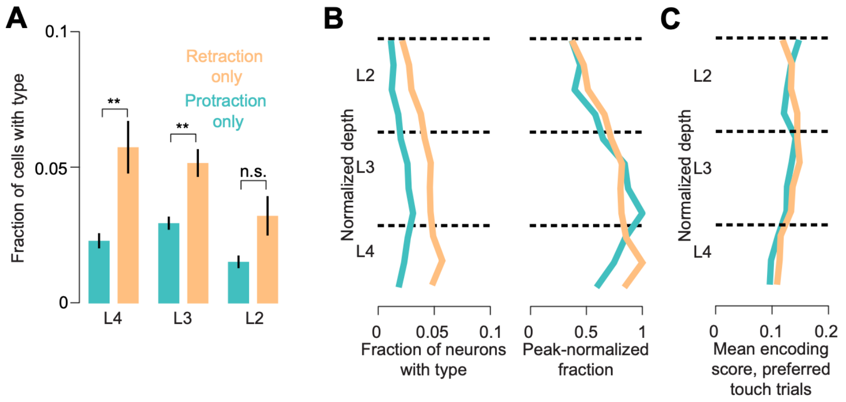

**Figure S6, Related to Figure 3. Laminar distribution of unidirectional single-whisker cells by directional preference.**

**A)** Frequency of protraction and retraction preferring unidirectional single-whisker cells for L4, L3, and L2. Bars show mean ± S.E.M. (n=7 mice).

**B)** Distribution of directional cell types as a function of normalized laminar depth (Methods). Left, fraction of cells at a given laminar depth. Right, Normalized fraction.

**C)** Encoding score for given directional cell type as a function of normalized depth. Encoding score was calculated only for trials of the preferred type.

#### Voelcker and Peron, Figure S7

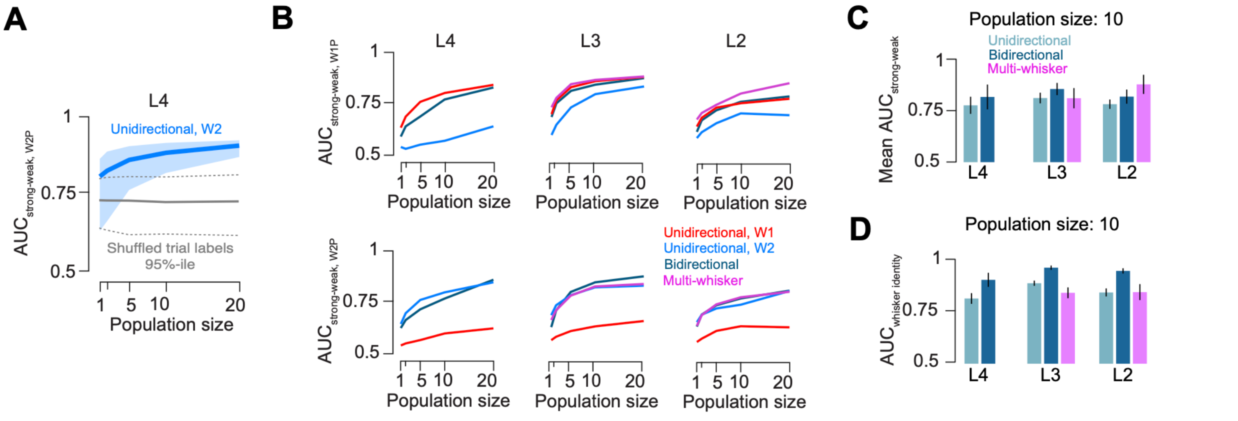

**Figure S7, Related to Figure 5. Population decoding of touch features.**

**A)** Population decoding of strong vs. weak W2P touches in L4 of an example mouse for varying numbers of unidirectional whisker 2 neurons. Blue line, median; light blue shading, 2.5$^{th}$ to 97.5$^{th}$ percentile (n = 1,000 repetitions per population size; Methods). Grey line, same analysis but with shuffled trial labels (Methods). Stippled grey line, 2.5$^{th}$ and 97.5$^{th}$ percentile of shuffled distribution.

**B)** Decoding as a function of population size for strong vs. weak W1P (top) and W2P (bottom) touches using different layers and cell types. Dark lines are means across all mice (n=7).

**C)** Decoding of strong vs. weak touches across all four single whisker touch types, shown for 10 neuron populations across cell types and layers. Symbols, individual mice. Bars indicate mean ± S.E.M. (n=7 mice).

**D)** Decoding of whisker 1 vs. whisker 2 touches for 10 neuron populations.
